# Inhibiting the prostaglandin transporter PGT induces non-canonical thermogenesis at thermoneutrality

**DOI:** 10.1101/836288

**Authors:** Victor J Pai, Run Lu, Licheng Wu, Marina Garcia Macia, Wade R Koba, Yuling Chi, Rajat Singh, Gary J Schwartz, Victor L Schuster

## Abstract

Prostaglandins play fundamental roles in adipose tissue function. While prostaglandin F_2α_ inhibits adipogenesis, prostaglandin E_2_ promotes adipose beiging. PGF_2α_ and PGE_2_ are both inactivated through uptake by the plasma membrane transporter (PGT). We hypothesized that inhibiting PGT would increase PGF_2α_ and PGE_2_ levels, thereby reducing white fat expansion and inducing beiging. Consistent with this hypothesis, inhibiting PGT in mice on high fat diet via genetic knockout or pharmacological blockade reduced body fat stores and induced thermogenesis at thermoneutrality. Inguinal white adipose tissue (iWAT) of these mice exhibited robust UCP1-independent thermogenesis characterized by mitochondrial expansion, coupling of O_2_ consumption to ATP synthesis, and induction of the creatine pathway. Enhanced coupled respiration persisted in PGT-KO iWAT adipocytes in a creatine shuttle-dependent manner. Thus, inhibiting PGT increases mitochondrial biogenesis and coupled respiration—each supported by the creatine pathway in a system lacking UCP1 expression—revealing PGT as a promising drug target against obesity.

## INTRODUCTION

Prostaglandins are 20-carbon fatty acid signalling molecules that are released by diverse cells types, including adipocytes ^1–3^. PGF_2α_ and PGE_2_ bind to their respective cell surface G protein-coupled receptors and activate a variety of downstream signalling events. Although PGF_2α_ and PGE_2_ are stable in plasma at 37°C, they do not function as circulating hormones. Rather, they are taken up by the broadly expressed prostaglandin reuptake carrier PGT (SLCO2a1) and delivered to a cytoplasmic oxidase for enzymatic oxidative inactivation ^4, 5^. PGT is the rate-limiting step in this two-step inactivation ^6^. The affinities of PGF_2α_ and PGE_2_ for their cognate receptors and for PGT are similar; because they compete for ligand, altering the rate of PGF_2α_ and PGE_2_ uptake by PGT reciprocally alters the degree of receptor signalling ^7^.

PGF_2α_ and PGE_2_ modulate adipose biology. In white adipose tissue (WAT), PGF_2α_ binds to its Gq-coupled receptor (FP) on adipocyte precursor cells, inhibiting adipocyte differentiation and lipogenesis ^8^. This effect is evident in humans when topical therapeutic PGF_2α_ analogues shrink the size of periorbital fat pads ^9^. Conversely, mice lacking PGF_2α_ synthase exhibit increased body fat on both normal and high fat diets ^10^.

In contrast, PGE_2_ appears to play a role primarily in inducing beige fat. When cold exposure stimulates sympathetic nervous outflow, the resulting activation of adipocyte adrenergic receptors by norepinephrine stimulates white adipocytes to synthesize PGE_2_. The latter enhances beige conversion and expression of uncoupling protein 1 (UCP1), thereby amplifying the cold response ^11–14^.

Based on these effects of PGF_2α_ and PGE_2_ on adipocyte biology, we hypothesized that genetically deleting PGT globally in mice (“PGT-KO”) would increase both systemic PGE_2_ and PGF_2α_, resulting in UCP1 induction and a lean phenotype. We found that, although PGT-KO mice are lean due to beige transformation of iWAT and thermogenesis, these processes are induced at thermoneutrality and do not require UCP1. Indeed, UCP1 knockout mice develop comparable iWAT beige transformation and thermogenesis when PGT is blocked pharmacologically. Suppression of UCP1 in PG-KO mice is secondary to suppression of PPARγ by FP receptor activation. The findings suggest that targeting PGT therapeutically may offer a novel approach for inducing a lean phenotype without UCP1.

## RESULTS

### PGT global knockout mice (PGT-KO mice) exhibit a lean phenotype

Results from both male and female KO mice are presented without stratification by sex because mice of both sexes exhibited the same metabolic phenotype. PGT-KO mice had elevated urinary PGE_2_ and PGF_2α_ excretion rates, indicating impaired systemic prostaglandin metabolism (Figure 1A). PGT-KO mice exhibited reductions in waist circumference (Figure 1 B-D), subcutaneous white adipose tissue (WAT) (Figure 1 E-F), visceral (gonadal) white adipose tissue (gWAT) (Figure 1 G-I), dermal fat (Figure 1 J-L), liver steatosis (Figure 1 M,N), and whole-body fat by echo-MRI composition analysis (Figure 1 O). PGT was differentially expressed in gWAT, iWAT, and interscapular brown adipose tissue (iBAT); compared to WT mice, the masses of these three fat depots in PGT-KO mice were reduced proportional to PGT expression (Supplementary Figure 1). PGT-KO mice displayed improved glucose tolerance compared to controls (Figures 1P and Supplementary Figure 1). WAT leptin gene expression, fasting serum leptin, and fasting serum free fatty acids were reduced in PGT-KO mice, whereas fasting serum adiponectin and insulin concentrations were unchanged (Supplementary Figure 1). Histology revealed that adipocytes of PGT-KO iWAT and iBAT depots were smaller than those of WT mice, and that PGT-KO iWAT contained multilocular adipocytes (Supplementary Figure 1).

**Figure 1.**
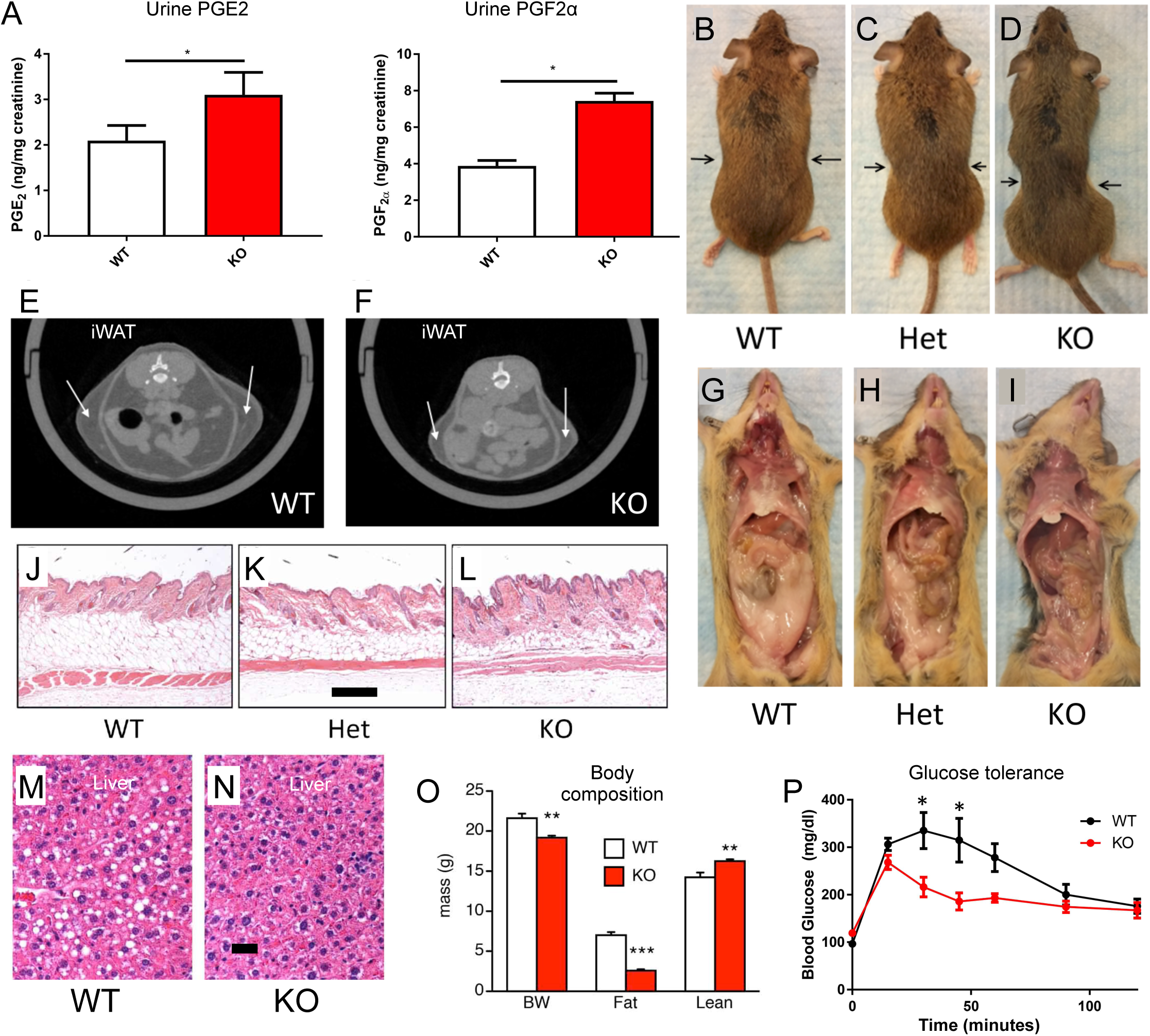
PGT-KO mice exhibit a lean phenotype. **(A)** Increased urinary concentrations of PGE_2_ and PGF_2α_ in PGT-KO mice, n=4 per group. **(B-D)** Representative gene-dosing effect on waist circumference of WT, PGT heterozygote, and PGT-KO mice. **(E-F)** Representative CT images of PGT WT and KO mice. Arrows indicate subcutaneous white adipose tissue (iWAT). **(G-I)** Representative visceral adipose tissue in WT, PGT heterozygote, and PGT-KO mice. **(J-L)** H&E sections of dermal adipose tissue in WT, PGT heterozygote, and PGT-KO mice. Bar = 50 µm. **(M-N)** Representative H&E sections of liver in WT and PGT-KO mice. Bar = 10 µm. **(O)** Quantification of total body lean and fat mass by echoMRI, n=4 per group. **(P)** Glucose tolerance test of WT and PGT-KO mice. N=4 per group. All mice housed at ambient temperature and fed a 9% fat diet by weight. Values are mean ± SEM. (*P<0.05, **P<0.01, ***P<0.001, versus respective control; Student’s t-test). For B-D, E-F, G-I, J-L, and M-N example shown were littermates.

### PGT-KO mice display increased energy expenditure due to beige induction in the iWAT depot

Although PGT-KO mice exhibited a 2-fold increase in food intake (Figures 2A and Supplementary Figure 2), there was no difference in stool weight, stool fatty acids, or intestinal integrity (Supplementary Figure 2), indicating that neither reduced energy intake nor malabsorption was not the cause of the lean phenotype. Because PGT-KO mice are lean despite a higher energy intake, they must be dissipating the excess energy as work and/or heat ^15^. Infrared beam interruption assay revealed an increase only of Y axis activity in PGT-KO mice, which was clustered at the onset of the active phase, a pattern indicative of hunger (Figure 2B) ^16^. PGT-KO mice exhibited an increase in O_2_ consumption (VO_2_) per lean body mass by indirect calorimetry (Figure 2C); under these experimental conditions, the observed change in VO_2_ cannot be attributed to the small increase in activity ^17^. To assess which tissues account for the whole-animal increase in thermogenesis, we injected mice with tracer deoxyglucose (F-18 FDG) and harvested tissues for analysis of uptake. Neither skeletal muscle nor interscapular brown adipose tissue (iBAT) displayed increased glucose uptake (Figure 2D). Further examination of skeletal muscle revealed no evidence for mitochondrial expansion or enhanced VO_2_ (Supplementary Figure 2). In contrast, iWAT exhibited a significant increase in F-18 FDG uptake (Figure 2D). Moreover, iWAT tissue explants from PGT-KO mice appeared visually “browned” (Supplementary Figure 2). PGT-KO iWAT exhibited induction of mitochondrial citrate synthase activity (Figure 2E), of browning genes (except UCP1) (Figure 2F), and of VO_2_ (Figure 2G). Extrapolating citrate synthase activity and VO_2_ of iWAT explants to the entire iWAT fat pad, or to the whole mouse, revealed a significant thermogenic capacity of this depot in PGT-KO mice (Figure 2E,G), a finding in agreement with the F-18 FDG results. Comparable extrapolations using liver citrate synthase data were unremarkable (Supplementary Figure 2).

**Figure 2.**
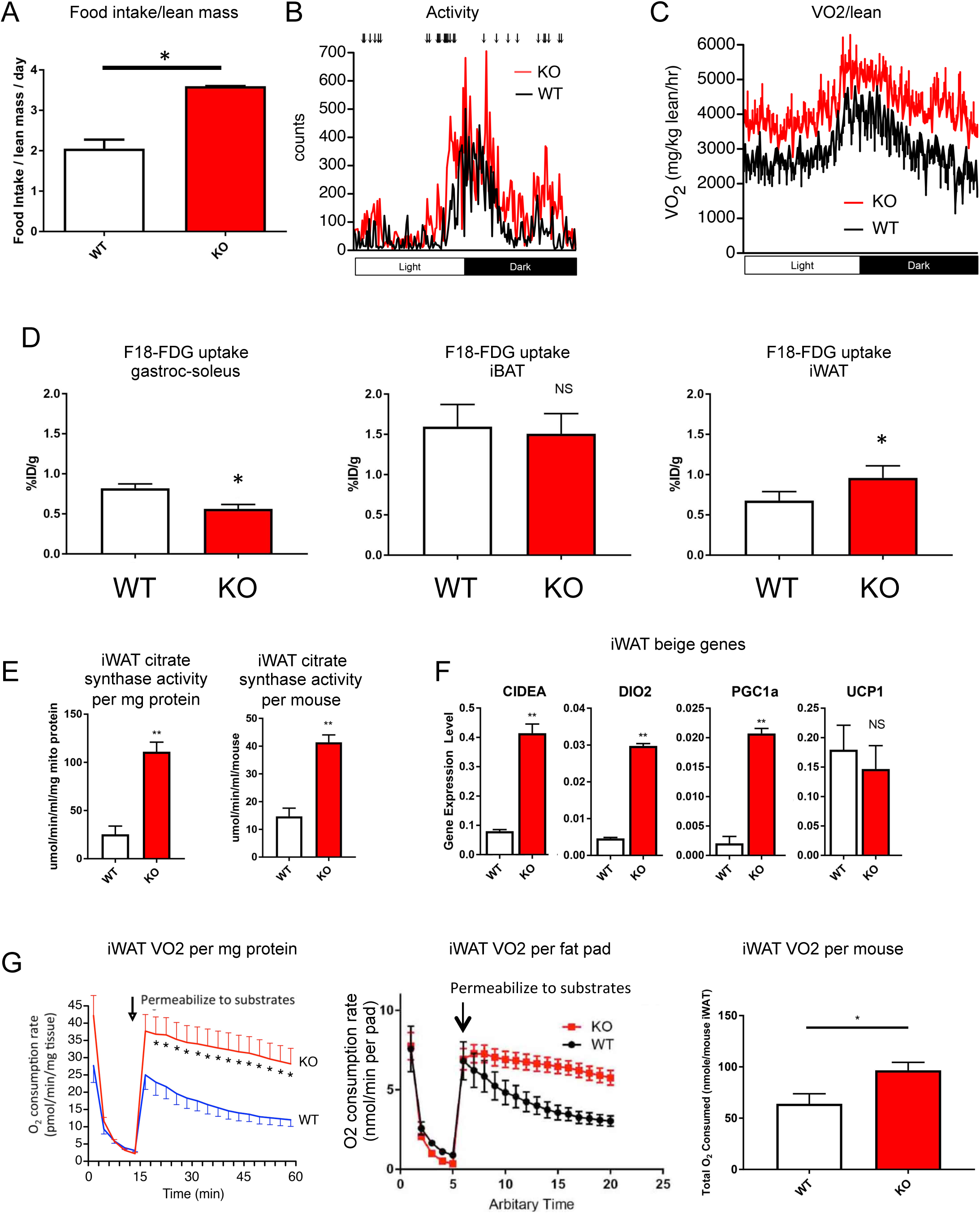
PGT-KO mice display increased energy expenditure with beige induction in the iWAT depot. **(A)** Increased food intake in PGT-KO mice, n=4 per group. **(B)** WT and PGT-KO mouse activity over 24 hours as measured by infrared beam break. Arrows indicate time points at which mean activity in PGT-KO mice is significantly greater (p<0.05) than that of WT mice. **(C)** Increase in VO_2_ per lean body mass in PGT-KO mice as measured by indirect calorimetry, n=4 per group. **(D)** 18-fluorodeoxyglucose uptake by gastroc-soleus skeletal muscle, interscapular brown adipose tissue (iBAT), and iWAT, n=4 per group. Data represent total uptake for the entire designated tissue of the mouse. **(E)** Increased citrate synthase activity in isolated PGT-KO iWAT mitochondria, n=4 per group, as activity per mg protein and extrapolated to whole mouse iWAT using depot weights. **(F)** Gene expression analysis of browning gene markers Cidea, Dio2, PGC1α, and UCP1 in iWAT of WT and PGT-KO mice by qRT-PCR, n=8 per group. **(G)** Increased oxygen consumption rate (OCR) in iWAT, measured by Seahorse assay, expressed as OCR per minute per mg tissue, extrapolated to entire iWAT fat pad by weight, and extrapolated to the entire mouse based on the weight of both iWAT fat pads. Cell membranes were permeabilized to substrates with digitonin as indicated, n=10. All mice housed at ambient temperature and fed a 9% fat diet by weight. Values are mean ± SEM. (*P<0.05, **P<0.01, versus respective control; Student’s t-test)

### WAT beige induction in PGT-KO mice represents “primary browning”

Skin or tail disorders in mice housed at ambient temperature can cause heat loss, resulting in “secondary browning” of WAT ^18^. To address secondary browning as phenotypic driver in PGT-KO mice, animals were tested in a water repulsion assay that detects heat loss from skin disorders ^19^. PGT-KO mice retained less water after immersion and defended body temperature as well as control mice (Supplementary Figure 3). To further exclude secondary browning, we constructed a thermal preference assay in which mice can choose freely amongst cages held at 22°C, 27°C, or 32°C (Supplementary Figure 3) ^20^. We validated the assay by determining the shift in thermal preference of C57BL/6J mice before and after depilation, which induces heat loss ^21^; fur removal induced a large shift in preference to 32°C (Supplementary Figure 3). PGT wild type mice housed at 22°C and then assessed over 24 hours in the preference assay demonstrated an integrated preference for the 32°C cage, whereas similarly housed and assayed PGT-KO mice displayed a preference distribution that was shifted toward cooler cages; these behaviours were more pronounced during the inactive (light) phase than the active (dark) phase (Figure 3A). Housing mice at 32°C before subjecting them to the thermal preference assay shifted all mice toward a warmer preference in the assay compared to those housed at 22°C; nonetheless, the population time budget distribution for PGT-KO mice compared to wild type controls remained shifted overall toward a cooler preference (Figure 3B).

**Figure 3.**
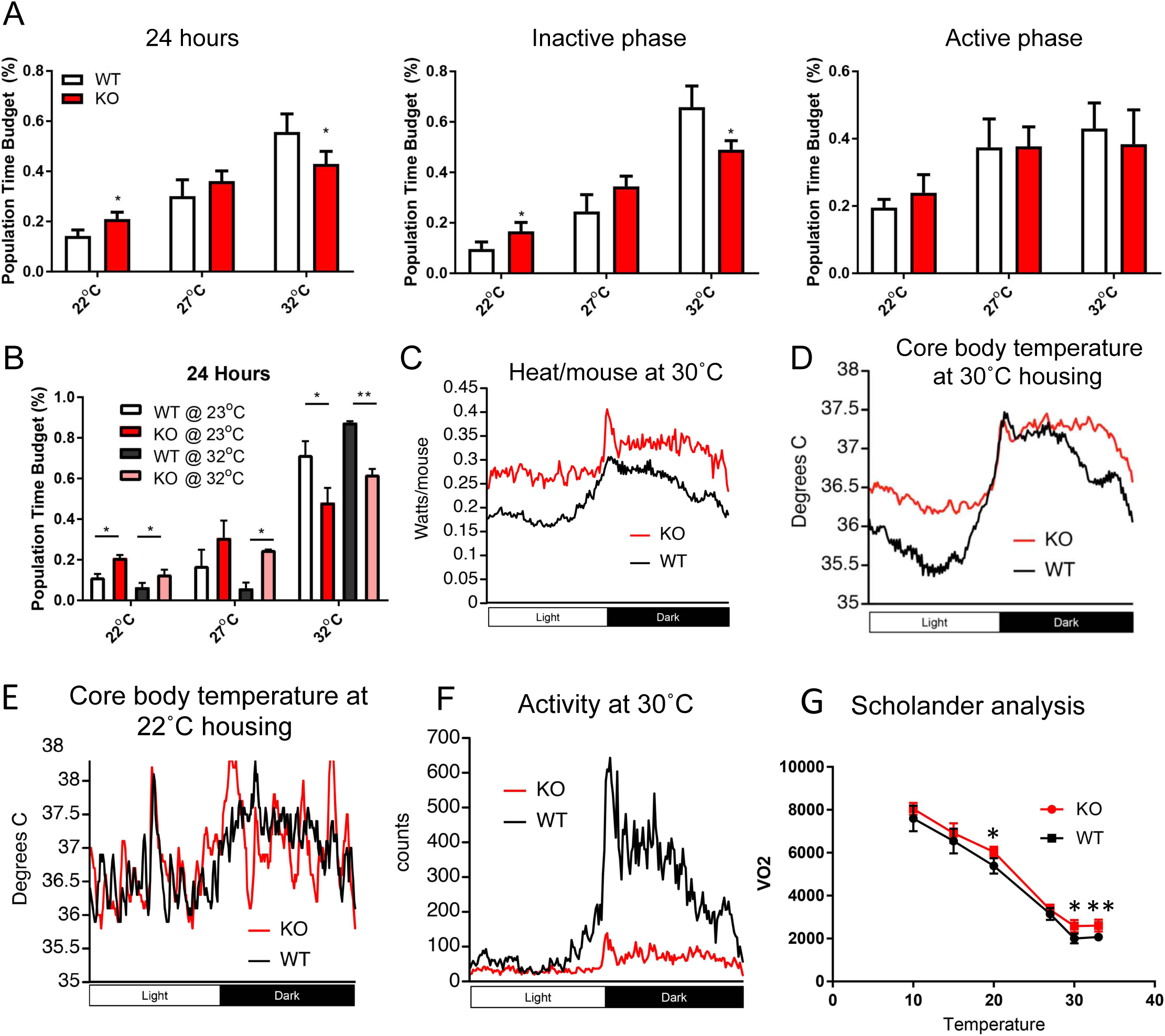
Primary thermogenesis in PGT-KO mice. **(A)** Thermopreference assay of WT and PGT-KO mice over the 24 hour diurnal cycle (left panel), and displayed separately in the inactive and active phases (middle and right panels), n=6. **(B)** Shifting of thermopreference in WT and PGT-KO mice after acclimation to either ambient temperature or to thermoneutrality, n=4 per group. **(C)** Increased heat generation in PGT-KO mice as measured by indirect calorimetry, n=4 per group. **(D)** Increased core body temperature in PGT-KO mice at 30°C, as measured by intraperitoneal probe, n=4 per group. **(E)** Normal core body temperature in PGT-KO mice at 22°C, as measured by intraperitoneal probe, n=4 per group. **(F)** Decreased activity of PGT-KO mice housed at 30°C, as measured by infrared beam break, n=4 per group. **(G)** Scholander plot analysis of WT and PGT-KO mice, n=8 per group. For (A), mice were housed at ambient temperature. For (B), the housing acclimation temperature prior to the acute thermopreference assay is shown in the inset. For (C-G), mice were housed at 30°C for ≥ 1 month before the respective assay. In all cases, mice were eating 9% fat diet by weight. Values are mean ± SEM. (*P<0.05, **P<0.01 versus respective control; Student’s t-test with Bonferroni correction where applicable.).

As with mice housed at 22°C, PGT-KO mice housed for 1 month at thermoneutrality (30°C) displayed increased thermogenesis compared to control mice (Figures 3C and Supplementary Figure 3). Core body temperature of PGT-KO mice housed at 30°C was higher than that of control mice during the inactive and late active phases (Figure 3D). In contrast, core body temperature of PGT-KO mice housed at 22°C was not different from that of control mice (Figure 3E). That PGT-KO mice dissipate their incremental heat when housed at 22°C, but not at 30°C, indicates that they have hyperthermia, rather than fever ^18^. In accord with hyperthermia, PGT-KO mice housed at 30°C displayed reduced spontaneous activity compared to mice housed at 22°C (compare Figure 3F to Figure 2B). Scholander analysis ^22^ failed to indicate heat loss in PGT-KO mice, i.e. there was no differential increase in VO_2_ of PGT-KO mice upon reducing environmental temperature ^23^; rather, PGT-KO mice exhibited an increase in VO_2_ only at thermoneutrality (Figure 3G). To test further the hypothesis that PGT-KO mice housed at 30°C are thermogenic, we transferred wild type control and PGT-KO mice from 30°C housing acutely to 4°C. Control mice defended core body temperature poorly and engaged in shivering, as determined by leak of muscle creatine kinase ^24^, whereas PGT-KO mice were able to defend body temperature with no apparent shivering (Supplementary Figure 3).

### WAT beige induction in PGT-KO mice persists on high-fat diet

To render these results more translatable to human obesity ^25^, we fed mice housed at 30°C a 60% high fat diet (HFD) for 1 month. The lean phenotype persisted under these conditions, with a reduction in body weight and total body fat and an increase in VO_2_ per lean body mass (Supplementary Figure 3). Explants of iWAT from PGT-KO mice revealed increased O2 consumption compared to controls (Supplementary Figure 3). Finally, analysis of WAT gene expression revealed an increase in brown and beige markers in iWAT, but not gWAT, of PGT-KO mice housed at 30°C on HFD (Supplementary Figure 3).

### Inhibiting PGT pharmacologically reproduces the knockout phenotype

To avoid possible confounding effects of altered adipose development when PGT is deleted from the single cell stage onward, as in PGT-KO mice, and to test whether inhibiting PGT on a pure C57BL/6 genetic background (as opposed to a mixed 129/BL6 genetic background) also results in thermogenesis ^26^, we administered a high-affinity PGT inhibitor ^27^ intraperitoneally to 2 month old C57BL/6J mice for 80-90 days. This phenocopied the results in PGT-KO mice as well as our previous results using a lower-affinity PGT inhibitor ^28^, producing an increase in urinary PGE_2_ and PGF_2α_ excretion (compare Figure 1A and Supplementary Figure 4). In C57BL/6J mice consuming a high fat diet, pharmacological PGT inhibition caused no change in food intake, but reduced body weight gain, a change that was entirely attributable to reduced fat accretion (Supplementary Figure 4). Inhibitor-treated mice exhibited higher O_2_ consumption rate as well as improved glucose disposal compared to vehicle-treated controls (Supplementary Figure 4). Finally, the PGT inhibitor caused induction of the beige genes Dio2 and Cidea (Supplementary Figure 4).

### Thermogenesis induced by deleting PGT is independent of UCP1

The data presented so far indicate that both PGT deletion and pharmacological PGT inhibition induce browning and thermogenesis of iWAT (Figures 2 and Supplementary Figure 4). If this thermogenesis is utilizing the canonical, UCP1-mediated pathway of uncoupled respiration, then gene expression levels of UCP1 in iWAT of PGT-KO mice should be increased over WT. However, UCP1 gene expression in this depot was not elevated in PGT-KO mice housed either at 22°C (Figure 2F) or at 30°C (Figure 4A). Indeed, in mice exposed to 4°C acutely for 15 hours, UCP1 gene expression in PGT-KO iWAT was suppressed relative to that of WT controls (Figure 4B). The lack of engagement of UCP1 in iWAT and iBAT of PGT-KO mice can be appreciated from UCP1 Q-PCR Ct values in these depots; the Ct’s were numerically higher (indicating lower mRNA expression) in iWAT and iBAT of PGT-KO mice housed at 22°C and 30°C, and in iWAT of mice exposed to 4°C for 16 hours; the only exception was iBAT of PGT-KO mice after 4°C exposure (Supplementary Figure 5).

**Figure 4.**
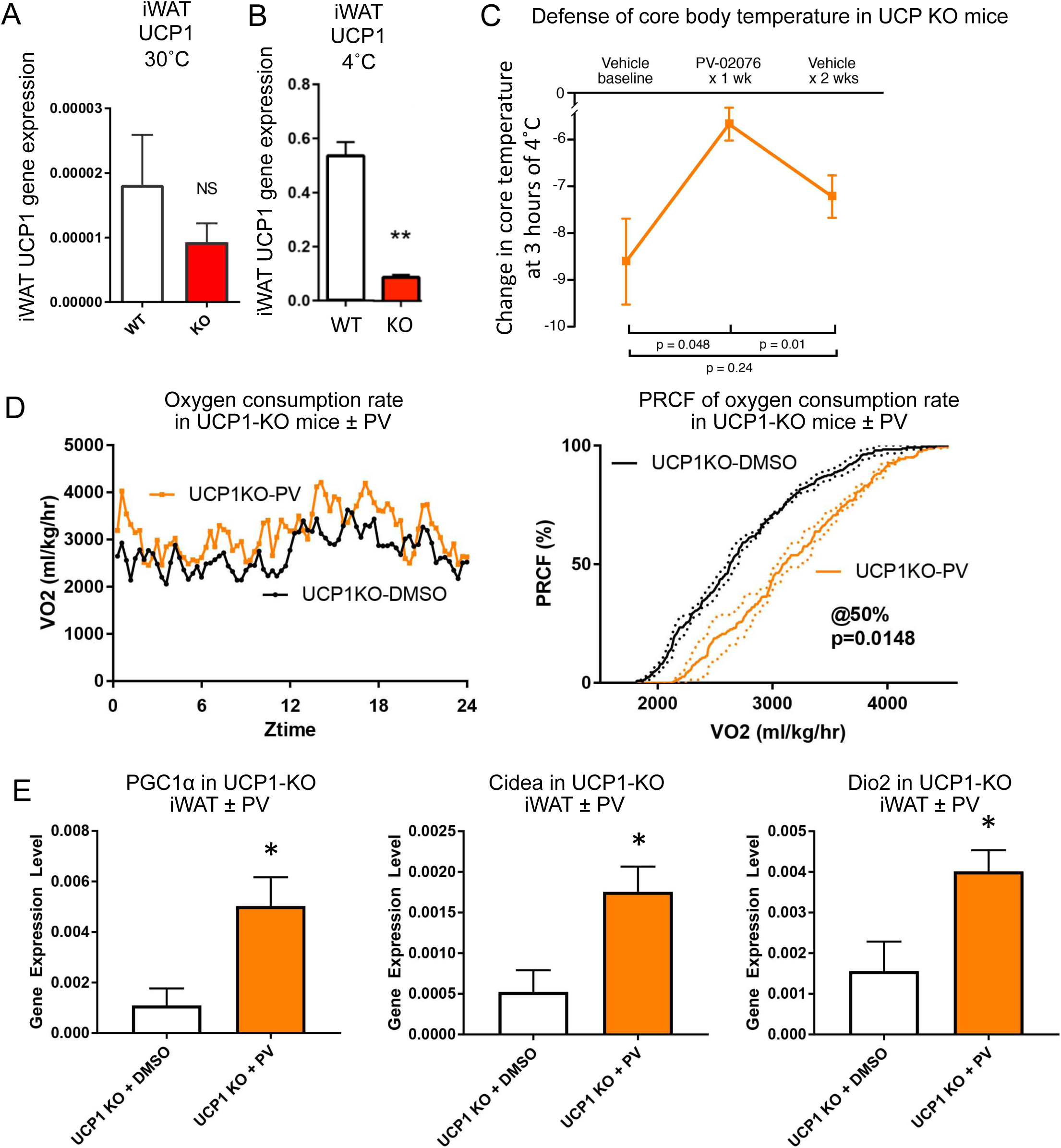
PGT deletion or pharmacological inhibition induces UCP1-independent thermogenesis. **(A-B)** iWAT UCP1 gene expression in PGT-KO mice housed at thermoneutrality (A) and after 16 hours’ exposure to 4°C (B). **(C)** Decrease in core body temperature upon 2 hours’ acute exposure to 4°C in UCP1-KO mice given vehicle for 1 week (left); the same mice after receiving PGT inhibitor PV-02076 for 1 week (centre); and the same mice after inhibitor washout (vehicle) for 2 weeks (right). n=6. **(D)** UCP1-KO mice exhibit Increase in VO_2_ when given PGT inhibitor PV-02076 (left) compared to DMSO control. VO_2_ data as percent relative cumulative frequency (PRCF) analysis ^75^ (right), presented as mean ± SEM, n=4 per group. At a PRCF of 50%, the DMSO mean VO2 = 2636± 35 and the PV mean VO2 = 3030±78, p = 0.015 by Student’s t-test. **(E)** Induction of browning gene expression markers in UCP1-KO mice given vehicle or PV-02076, n=4 per group. All mice were housed at thermoneutrality except for cold exposure in (C). Values are mean ± SEM. (*P<0.05, **P<0.01, versus respective control; Student’s t-test (A-B, D-E) and one way ANOVA (C).

To explore further the concept that inhibiting PGT induces thermogenesis in the absence of UCP1 induction, we assessed the defence of core body temperature in UCP1 knockout mice (UCP1-KO) ^29^. We housed UCP1-KO mice ^30^ at 30°C, administered vehicle (DMSO) alone, and brought them acutely to 4°C, whereupon they defended core body temperature poorly, exhibiting a mean drop of core body temperature of ∼9°C over 3 hours (Figure 4C). We then administered the PGT inhibitor for 7 days and repeated the assay; the same mice now exhibited improved acute defence of core body temperature, with a mean drop in core body temperature of < 6°C at 3 hours (Figure 4C). Finally, we washed out the inhibitor for 2 weeks; the same mice reverted toward their previous state of impaired defence of core body temperature (Figure 4C). Separately, we housed UCP1-KO mice at 30°C and treated them with vehicle or PGT inhibitor. The inhibitor induced thermogenesis, as judged by indirect calorimetry and induction of the beige genes PGC1α, Cidea, and Dio2 in iWAT (Figure 4D-E). Thus, inhibiting PGT at thermoneutrality induces iWAT-based thermogenesis in the complete absence of UCP1.

### Factors contributing to suppression of iWAT UCP1 gene expression

We explored two possible mechanisms for suppression of UCP1 in PGT-KO mice. First, because PGE_2_ is known to act through inhibitory EP_3_ receptors on sympathetic nerve endings to reduce norepinephrine release ^31–33^, and because PGE_2_ is elevated in PGT-KO mice (Figure 1A) and in PGT inhibitor-treated mice (Supplementary Figure 4), we measured urinary norepinephrine, an index of systemic norepinephrine release from sympathetic nerve terminals ^34–36^. As shown in Figure 5A, urinary norepinephrine excretion was markedly reduced, both in PGT-KO mice housed at 30°C and in control mice administered the PGT inhibitor PV and housed at 22°C. As a functional correlate of these measurements, we also measured systolic and diastolic arterial blood pressure and heart rate in vehicle- and inhibitor-treated mice. As shown in Supplementary Figure 5, although the PGT inhibitor produced no change in blood pressure in these normotensive mice, a finding in accord with our previous report ^37^, pharmacological PGT inhibition lowered resting heart rate significantly, an indicator of reduced sympathetic tone ^38, 39^. In contrast to norepinephrine, urinary epinephrine, an index of systemic epinephrine release from the adrenal medulla ^35^, was not affected, nor was the expression in iWAT of tyrosine hydroxylase, the rate limiting step in catecholamine synthesis (Supplementary Figure 5).

**Figure 5.**
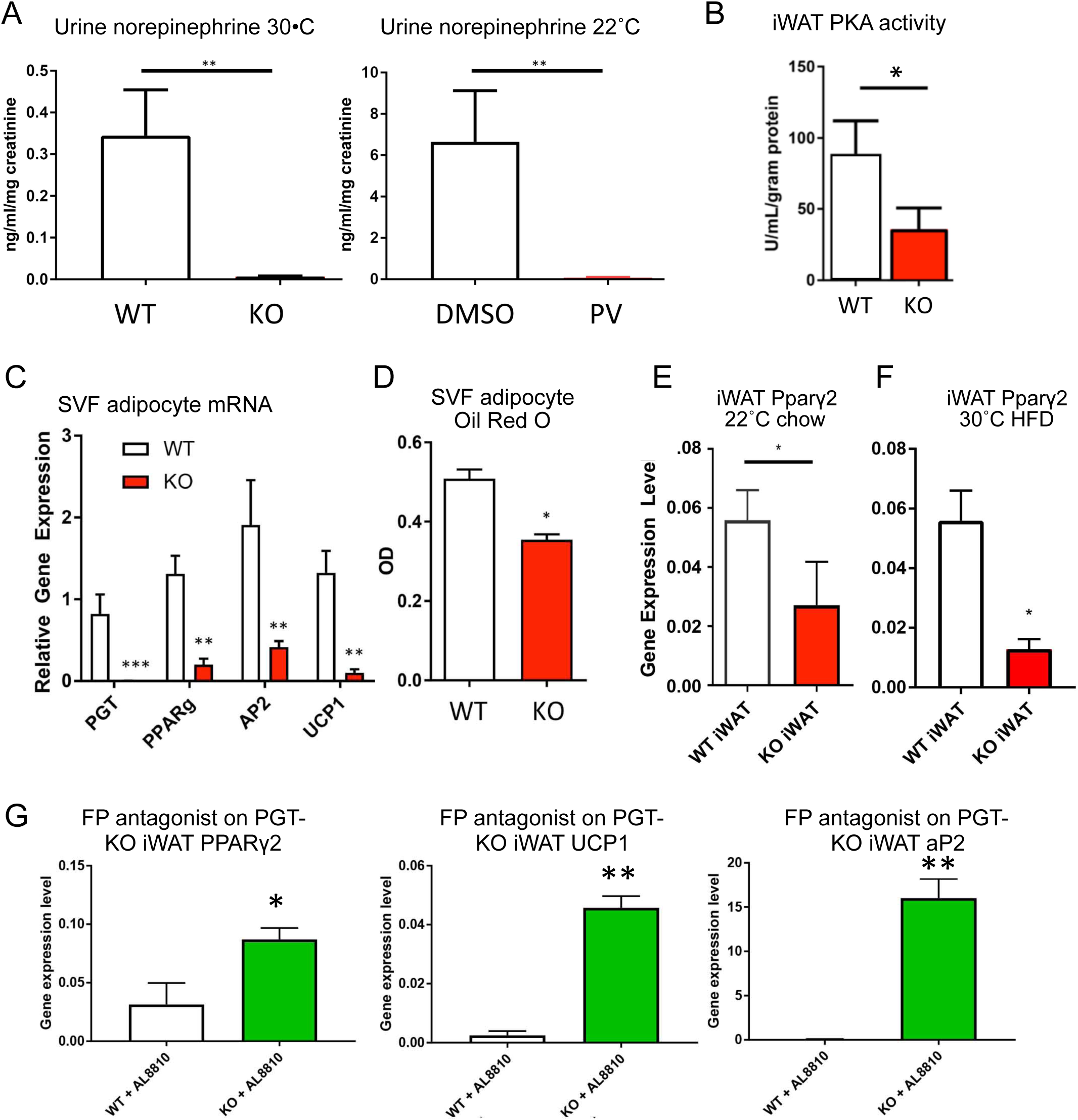
Mechanisms of suppression of UCP1 in PGT-KO mice. **(A)** Inhibition of systemic norepinephrine release in PGT-KO mice housed at 30°C (n=4 per group) and in C57BL/6J mice given PV-02076 for 1 month at 22°C (n=8 per group), as measured by urinary norepinephrine levels. **(B)** Decreased protein kinase A activity in iWAT of PGT-KO mice. **(C)** Decreased expression of adipocyte markers in adipocytes derived from stromal vascular fraction (SVF) of WT and PGT-KO iWAT as measured by qRT-PCR **(D)** Decreased lipid droplet accumulation in SVF-derived adipocytes as measured by oil red O staining. **(E)** Decreased PPARγ2 expression in adipocytes induced *in vitro* from SVFs derived from iWAT of WT and PGT-KO mice housed at 22°C eating 9% fat diet, and from comparable mice housed at 30°C and eating a 60% high fat diet. For figures (C-D), values are mean ± SEM of at least 3 independent experiments. **(G)** Rescue of PPARγ2, UCP1, and aP2 gene expression in iWAT of PGT-KO mice given FP antagonist AL-8810, as measured by qRT-PCR, n=4 per group. Values are mean ± SEM. (*P<0.05, **P<0.01, ***P<0.001, versus respective control; Student’s t-test)

Despite the loss of norepinephrine as a cyclic AMP agonist, protein kinase A (PKA) activity of PGT-KO iWAT exhibited only a modest reduction (Figure 5B). Because PKA in the iWAT depot of sympathectomized mice retains its ability to be activated by agonists ^40^, the persistent PKA activity seen here in PGT-KO iWAT suggests that chronically elevated PGE_2_ is functioning as a constitutive PKA activator ^12^ in lieu of the normal facultative adrenergic stimulus.

We also tested a second hypothesis, namely that suppression of iWAT UCP1 gene expression is intrinsic to the PGT-KO iWAT adipocyte. We isolated the stromal vascular fraction (SVF) from iWAT of PGT-KO mice and induced differentiation into adipocytes using standard stimuli ^41, 42^. Compared to adipocytes induced from wild type control mice, adipocytes induced from PGT-KO SVF displayed undetectable PGT expression, lower expression of a “white” adipocyte phenotype (reduced Oil Red O accumulation and aP2 gene expression), and suppressed expression of UCP1 and PPARγ (Figure 5C-D). The reduced PPARγ expression of *in vitro* adipocytes (Figure 5C) was confirmed in intact iWAT tissue from PGT-KO mice housed at both 22°C eating normal chow and in mice housed at 30°C eating high fat diet (Figure 5E-F). These data are consistent with a model in which the cAMP pathway in iWAT that is activated by PGE_2_ induces PGC1α expression, but the latter is incapable of increasing UCP1 transcription because its co-factor PPARγ is suppressed ^43–45^.

In considering the mechanism of iWAT PPARγ suppression, we noted reports that PGF_2α_, acting through its Gαq-coupled receptor FP, suppresses PPARγ, and hence UCP1, gene expression ^3, 8, 46^. To test the degree to which PGF_2α_ plays such a role in PGT-KO mice, we administered the specific FP receptor antagonist AL8810 ^47–49^ to WT and PGT-KO mice for 4 days. Figure 5G shows that blocking FP signalling reversed the effects of PGT-KO on iWAT PPARγ, UCP1, and aP2 gene expression, suggesting that the rise in PGF_2α_ from PGT-KO plays a dominant role in suppressing both iWAT UCP1 and the white adipocyte phenotype.

### Thermogenesis in PGT-KO iWAT is coupled to ATP synthesis and is associated with induction of the creatine shuttle pathway which, in turn, is dependent upon signalling through the PGF_2α_ receptor

Because UCP1 is not induced in PGT-KO iWAT, the incremental thermogenesis is unlikely to be uncoupled from ATP synthesis, as is the case with UCP1-derived thermogenesis. We examined this question directly by determining the O_2_ consumption rate of iWAT explants from WT and PGT-KO mice before and after inhibiting ATP synthase with oligomycin. These measurements revealed that the increment in iWAT O_2_ consumption is coupled to ATP synthesis (Figure 6A-B). In addition to the induction of iWAT browning genes in the absence of UCP1 ^50^, induction of elements of the classical creatine shuttle in this setting have also been reported ^24^. Here, iWAT of PGT-KO mice housed at thermoneutrality displayed induction of genes encoding the creatine transporter *Slc6a8 and* mitochondrial creatine kinases *Ckmt*1 and *Ckmt*2 (Figure 6C). Inhibiting PGT pharmacologically in C57BL/6 mice housed at 30°C induced Ckmt1 and Ckmt2 in iWAT (Figure 6D). In UCP1-KO mice housed under the same conditions, the PGT inhibitor induced iWAT expression of *Ckmt*1 and *Slc6a8* (Figure 6E), indicating that cold exposure of UCP1-KO mice is not required for induction of creatine shuttle components. To test the hypothesis that the creatine pathway contributes to whole-mouse thermogenesis in PGT-KO mice, we administered the *Slc6a8* transporter inhibitor β-guanidinopropionic acid (β-GPA) systemically as reported ^24^, however, we were unable to normalize the augmented VO_2_ of PGT-KO mice in this manner (Supplementary Figure 6).

**Figure 6.**
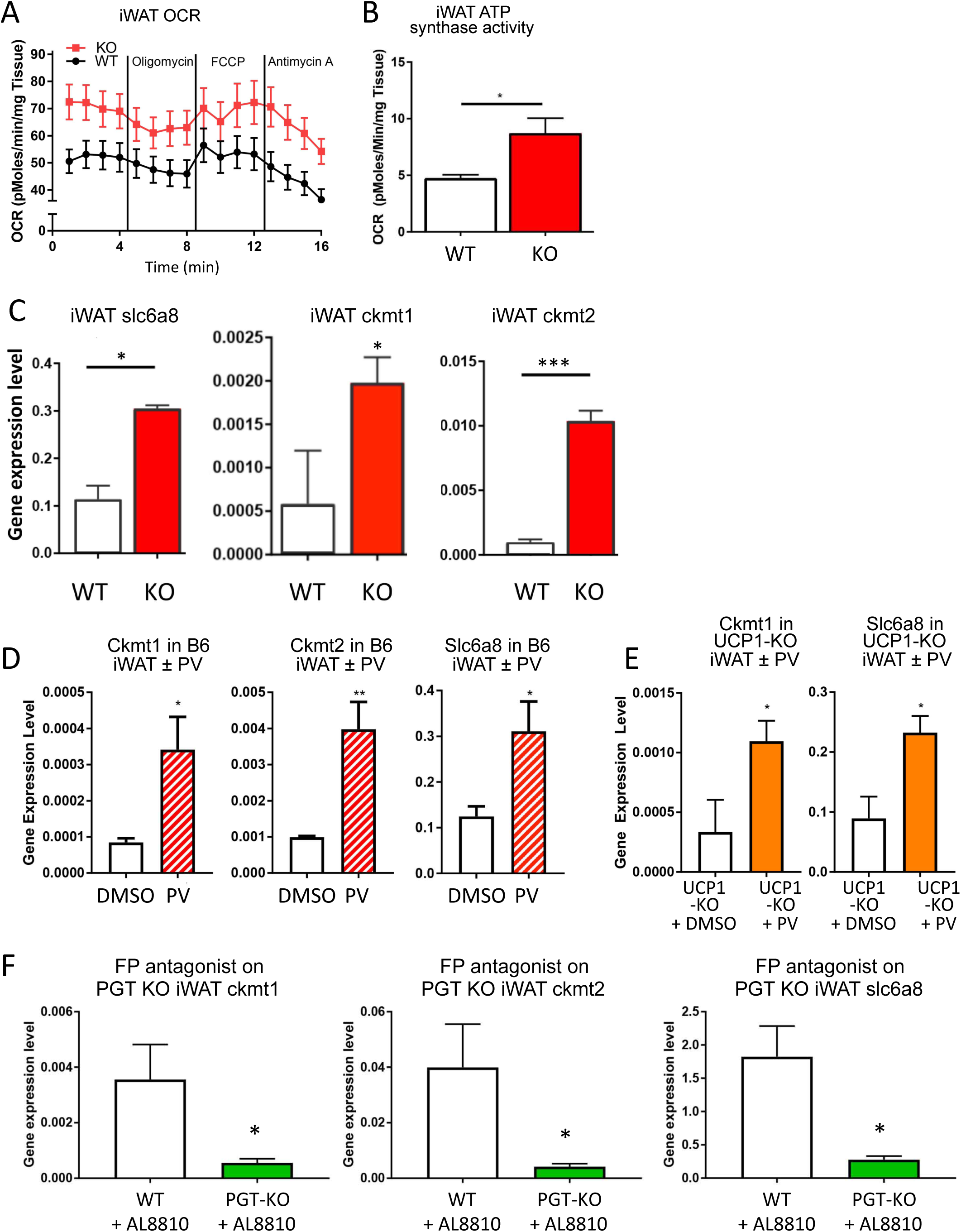
Increased ATP-coupled thermogenesis and creatine shuttle gene expression in PGT-KO mice. **(A-B)** Increased ATP synthase activity in iWAT of PGT-KO mice (n=8). ATP synthase activity is calculated as (average baseline OCR) – (average oligomycin OCR). **(C-F)** Creatine shuttle gene expression in iWAT: Ckmt = mitochondrial creatine kinase, Slc6a8 = Na^+^-creatine symporter. (C) iWAT of WT vs PGT-KO; (D) iWAT of C57BL6 mice administered vehicle (DMSO) or the PGT inhibitor PV-01076; (E) UCP1-KO mice administered DMSO or PV-02076 **(F)** Loss of induction of PGT-KO iWAT creatine shuttle genes after blockade of the PGF_2α_ receptor FP by AL8810. Values are mean ± SEM, (*P<0.05, **P<0.01, ***P<0.001, versus respective control; Student’s t-test). All mice housed at thermoneutrality and consuming 9% fat by weight diet.

Because the PGF_2α_ receptor FP plays a key role in suppressing UCP1 gene expression in PGT-KO mice (Figure 5G), we also explored the role of FP in control of the creatine shuttle pathway. Administering the FP antagonist AL8810 to PGT-KO mice reversed the induction of iWAT Ckmt1, Ckmt2, and Slc6a8 (Figure 6F). In addition to genes of the creatine shuttle, the muscle genes Serca1 (*Atp2b1)* and *Myf5* and were also induced in iWAT of PGT-KO mice, as were a number of genes of fatty acid β-oxidation (Supplementary Figure 6). In contrast, there was no induction of genes involved in either lipolysis or lipogenesis (Supplementary Figure 6).

### Thermogenesis in PGT-KO iWAT beige adipocytes is dependent upon creatine

To explore further the mechanism of thermogenesis in PGT-KO iWAT, we used adipocytes induced *in vitro* from the stromal vascular fraction. As with iWAT explants, adipocytes induced *in vitro* from PGT-KO iWAT exhibited an increase in coupled respiration (Figure 7 A-B) and induction of the creatinine shuttle gene Ckmt2 (Figure 7C). We validated β-GPA as an effective tool in *in vitro* by confirming its ability to inhibit O_2_ consumption in adipocytes derived from UCP1-KO mice ^24^ (Figure 7D). When applied to adipocytes derived from WT mice, β-GPA had no effect on O_2_ consumption, however, β-GPA returned the increased O_2_ consumption rate of PGT-KO adipocytes to control levels (Figure 7E), indicating a functional role for the creatine shuttle in the non-UCP1-mediated thermogenesis of PGT-KO iWAT beige adipocytes.

**Figure 7.**
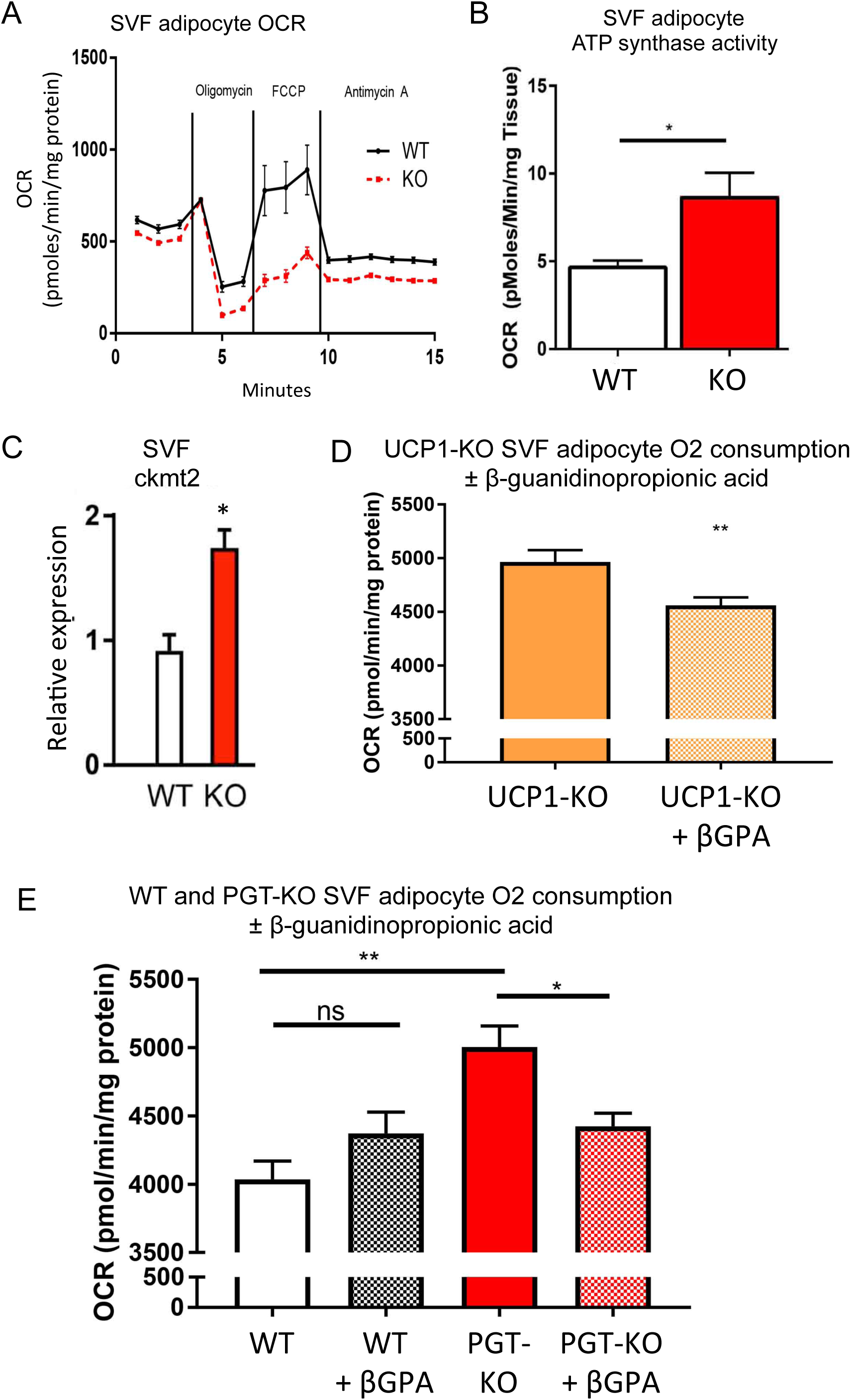
Non-canonical thermogenesis supported by the creatine shuttle in iWAT adipocytes in vitro. **(A)** Oxygen consumption rate (OCR) of SVF-derived adipocytes. **(B)** Oligomycin-sensitive OCR from A, equivalent to ATP synthase activity. **(C)** Up-regulation of Ckmt2 in PGT-KO adipocytes *in vitro*. **(D)** Inhibition of OCR by β-GPA in adipocytes *in vitro* derived from UCP1-KO iWAT. **(E)** Reversal *in vitro* by β-GPA of elevated OCR in adipocytes derived from PGT-KO iWAT. OCRs calculated as averages from 5 wells across 4 independent time points. n = 4. For (D) ** p < 0.01 by Student’s t-test; for (E) ** p < 0.01 by one-way ANOVA.

## DISCUSSION

The present studies demonstrate that genetically deleting, or pharmacologically inhibiting, the prostaglandin uptake carrier PGT in mice induces primary thermogenesis and reduced fat accretion in multiple adipose depots and in liver. Tissue-specific glucose uptake, O_2_ consumption, and gene expression changes indicate that the increased thermogenesis owes, at least in part, to beige transformation of subcutaneous inguinal white adipose tissue (iWAT). Thermogenesis in PGT-KO iWAT does not derive from canonical UCP1-based uncoupled respiration. Rather, UCP1 gene expression that might otherwise be stimulated by PGC-1α is, instead, suppressed due to activation of the PGF_2α_ receptor. The incremental respiration is coupled to ATP synthesis (“non-canonical thermogenesis”) and is accompanied by induction of the creatine shuttle pathway, which is functionally necessary for the increased respiration.

### Primary thermogenesis in PGT-KO mice

A compelling argument has been adduced that many cases of beige induction in mice housed at ambient temperatures are not primary, but rather are secondary due to heat loss through skin, fur, or tail ^18^. We addressed this issue in several complementary ways. First, PGT-KO mice exhibit normal water repulsion and a behavioural preference for a cooler, rather than warmer, environment, especially during the inactive phase. Second, Scholander curves on PGT-KO mice are inconsistent with heat loss. Third, PGT-KO mice housed at thermoneutrality maintain an increase of whole-mouse and iWAT O_2_ consumption, as well as induction of browning genes in iWAT. Fourth, the response of core body temperature in PGT-KO mice to changes in ambient temperature indicates that they have hyperthermia, not fever. Finally, the thermogenic pathway activated in PGT-KO mice at 30°C serves to defend PGT-KO mice from acute cold exposure, obviating the need to shiver. Taken together, the data argue strongly that iWAT beige induction in PGT-KO mice is primary and not secondary.

### Stimulation of mitochondriogenesis and coupled respiration

Recent studies have demonstrated that PGE_2_ plays an amplifying role in the so-called “canonical”, or UCP1-mediated, thermogenic response of WAT to cold exposure ^11, 12^. Specifically, norepinephrine increases cyclic AMP levels directly by activating β3-adrenergic receptors, and indirectly by inducing PGE_2_ synthesis that, itself, activates the same signalling cascade via EP_4_ receptors ^12^. By increasing systemic levels of PGE_2_ in PGT-KO mice, we hypothesized that the cyclic AMP pathway in mice housed under mild thermal stress (22°C) would be enhanced. Surprisingly, protein kinase A activity in iWAT of these PGT-KO mice was not increased, rather it was only moderately decreased. One possible explanation for this result is that systemic norepinephrine release, as determined by urinary excretion ^34, 35^, was markedly suppressed in PGT-KO mice and in mice administered a PGT inhibitor. This result is in accord with the known ability of PGE_2_ to suppress norepinephrine release from sympathetic nerve termini via EP_3_ receptors ^31–33^. Together, the data suggest that elevating PGE_2_ constitutively by blocking its metabolism directly stimulates the iWAT cAMP - protein kinase A pathway, while at the same time inhibiting facultative activation of this pathway by adrenergic agonists. The net result is constitutive activation of mitochondriogenesis, an increase in coupled respiration, and induction of genes of fatty acid β-oxidation.

### Constraints on UCP1 gene expression in PGT-KO iWAT

Whereas deleting PGT increased the expression of a broad array of iWAT browning genes in mice housed both at 30°C and 22°C, *Ucp1* gene expression was strongly suppressed, and could not be induced even by 16 hrs of exposure to 4°C. Although administering exogenous PGE_2_ alone to mice induces *Ucp1* in WAT ^12^, inhibiting PGT increases both PGF_2α_ and PGE_2_. In WAT, PGF_2α_ suppresses white adipocyte differentiation as well as *Ucp1* expression ^3, 8, 10, 51–53^. Both effects result from PGF_2α_ inhibiting PPARγ gene expression and function ^8, 54, 55^. In keeping with these known effects of PGF_2α_, PPARγ mRNA was reduced both in whole iWAT of PGT-KO mice and in adipocytes derived *in vitro* from PGT-KO iWAT, and blocking the PGF_2α_ receptor in PGT-KO mice rescued *Ucp1* gene expression. Experiments by Klepac et al indicate that directly stimulating the g protein Gαq, which is coupled to the PGF_2α_ receptor FP, also suppresses PPARγ and UCP1 ^46^. Because high concentrations of PGE_2_ can engage the EP_1_ receptor, which also signals through Gαq to suppress PPARγ ^3, 14^, it is possible that elevated levels of PGE_2_ may also have contributed to suppressing *Ucp1*. Taken together, the data are consistent with a model in which increased levels of PGE_2_ and PGF_2α_ stimulate iWAT expression of PGC1α, mitochondrial expansion, and expression of browning genes while simultaneously inhibiting expression of the PGC1α binding partner PPARγ, and thus UCP1 expression (Supplementary Figure 7). Despite the relatively low capacity of beige adipocyte mitochondria for ATP synthesis ^56^, the significant expansion of mitochondrial mass in iWAT of PGT-KO mice appears sufficient to support an increase in coupled O_2_ consumption. In this regard, it is noteworthy that both the Scholander curves and the modest increase in core body temperature at thermoneutrality, but not ambient, temperature of PGT-KO mice resemble those of voles bred for high aerobic capacity that have a 7% higher mass-adjusted basal metabolic rate compared to controls ^57, 58^.

### Role of genetic strain and thermoneutrality

Although UCP-KO mice on either a pure C57BL/6J or a pure 129/SvlmJ genetic background are markedly cold-sensitive, UCP1-KO mice on a mixed 129xBL/6 background are cold-resistant ^50, 59, 60^. Similarly, in the present studies, PGT-KO mice on a mixed 129xBL/6 background exhibited thermogenesis with improved cold tolerance despite suppression of UCP1. Importantly, by inhibiting PGT pharmacologically at thermoneutrality, thermogenesis and improved cold tolerance could be induced in mice on a pure C57BL/6J background, indicating that neither an F1 mixed genetic background nor cold exposure is required for the PGT inhibition effects. Although over-expressing cyclooxygenase-2 or adenosine monophosphate-activated protein kinase (AMPK) in mice housed at thermoneutrality has been reported to protect against diet-induced obesity ^11, 61, 62^, in the present model leanness, thermogenesis, and improved cold tolerance were all induced under thermoneutral conditions.

### Beige adipocyte types, creatine pathway, and cellular mechanisms responsible for UCP1-independent thermogenesis in PGT-KO iWAT

PGT mRNA in iWAT is expressed in an adipocyte precursor population ^63^. This cell specificity would position PGT for paracrine control ^7^ of white or beige adipogenesis. Further work is required to delineate both the target cell(s) of PGE_2_ and PGF_2α_ paracrine signalling in the iWAT depot, as well as the characteristics of the resulting beige adipocytes, especially since recent evidence suggests that a number of novel subtypes of beige adipocytes may exist ^24, 64–69^. To the extent that PGT-KO iWAT expresses components of the creatine shuttle, Myf5, and SERCA1, the corresponding thermogenic beige adipocytes may represent yet another novel cell type.

The inhibition of oxygen consumption by β-GPA in PGT-KO iWAT adipocytes *in vitro* is consistent with an emerging role of the creatine shuttle in beige adipocyte thermogenesis, especially in the absence of UCP1-mediated, uncoupled respiration ^24, 64, 65, 70^. Although the creatine pathway was initially identified in beige adipocytes derived from UCP1-KO mice, suggesting that UCP1 and the pathway vary reciprocally ^24^, the present results indicate that PGF_2α_ may independently regulate this pathway, at least in the absence of PGT. Thus, the PGF_2α_ receptor inhibitor simultaneously increased gene expression of UCP1 in PGT-KO iWAT (Figure 5G) while suppressing expression of creatine shuttle genes (Figure 6F). The lack of an effect of β-GPA on thermogenesis in intact mice (present study in PGT-KO mice, and ^61^) may reflect pharmacodynamic issues, or may indicate that the contribution of iWAT to overall thermogenesis in these models is less than that of other depots or tissues.

UCP1-independent thermogenesis by beige adipocytes requires activation of an alternative futile cycle ^71^. Candidates for the latter that have been put forward include uncoupling of sarcoendoplasmic reticulum calcium ATPase (SERCA) ^69^ and cycling of lipolysis-lipogenesis ^72^. The futile cycle(s) generating heat in iWAT of PGT-KO mice remain(s) to be identified.

The proposed model for PGT-KO iWAT is shown in Supplementary Figure 7. In this model, increased PGE_2_ in PGT-KO mice inhibits facultative norepinephrine release from sympathetic nerve terminals, PGE_2_ can still activate cAMP signalling constitutively to induce PGC1α activation and mitochondrial biogenesis. Increased β oxidation of fatty acids drives increased ATP synthesis by the expanded mitochondrial pool. The accompanying increase in PGF_2α_ in PGT-KO mice activates the receptor FP which, via Gαq signalling, reduces UCP1 gene expression by inhibiting PPARγ gene expression and induces components of the creatine shuttle. The increased ATP synthesis, via the creatine shuttle, supports UCP1-independent thermogenesis via (unidentified) futile cycle(s).

### Therapeutic implications

Although activating UCP1-mediated thermogenesis in humans obesity seems theoretically sound, in practice it has been difficult to activate UCP1 in subjects who are obese or beyond their young adult years, or to translate activation into meaningful weight loss in the target population ^73^. Instead, it has been argued that non-canonical thermogenesis is less efficient than UCP1-mediated thermogenesis ^74^, and therefore may be a preferable therapeutic pathway to target. The present results provide evidence that pharmacologically inhibiting PGT induces robust WAT non-canonical thermogenesis in mice housed at thermoneutrality and consuming a high fat diet, that is conditions mimicking those of obese human subjects. These findings raise PGT as a promising drug target against obesity.

## METHODS

### Animals

All animal procedures were performed under the guidelines of Albert Einstein College of Medicine’s Institutional Animal Care and Use Committee. The generation and rescue of PGT-KO mice were as reported previously ^76^. Animals were either housed at 22°C or 30°C, and fed either chow (5058, LabDiet, St. Louis, MO, USA) or high fat diet (D12492, Research Diets, New Brunswick, NJ, USA) depending on experimental conditions. PGT-KO mice are on a mixed 129/BL6 background. Wild type mice of the same genetic background were used as controls. For inhibitor studies, C57BL/6J wild type, and UCP1 knockout mice (B6.129-*Ucp1^tm1Kz^*/J), were obtained from Jackson Laboratory (Bar Harbor, ME).

The PGT inhibitor PV-02076 ^27^ was dissolved in DMSO and injected intraperitoneally at a dose of 20 mg/kg, with DMSO as a vehicle control. Mouse tissue was either fixed in 10% phosphate buffered formalin followed by 70% ethanol for sectioning and staining with hematoxylin and eosin, or was snap-frozen in liquid nitrogen and stored at −80°C. Mouse whole blood was collected by retro-orbital bleeding and allowed to coagulate at room temperature. Serum was collected by centrifuging whole blood samples at 5000 x g for 10 minutes. Samples were sent to the University of Cincinnati Mouse Metabolic Phenotype Centre, where serum free fatty acids, adiponectin, leptin levels were measured. Mouse liver triglyceride levels were measured by colorimetric assay according to manufacturer instructions (Cat# 10010303, Cayman Chemical, Ann Arbor, Michigan, USA). For food and water intake measurements, mice were housed individually, and food/water intake was measured daily for one week. Body fat composition was measured by echoMRI (Echo Medical Systems, Houston, Texas). CT studies were performed under isoflurane anaesthesia. Urine PGE_2_ and PGF_2α_ were measured by ELISA (Cayman Chemical); urine for these assays as well as stool for the assays below were collected by housing mice in metabolic cages over 2 weeks and samples were stored at −80°C. Urine creatinine was measured by LC-MS at the University of Alabama at Birmingham O’Brien Center Bioanalytical Core. Urine epinephrine and norepinephrine, collected from spot samples between 11:00 AM and 2:00 PM, were measured by ELISA (NBP2-62867, NOVUS Biologicals; BA E-6200, Rocky Mountain Diagnostics, Colorado Springs, Colorado, respectively) and normalized to urinary creatinine. Tissue protein kinase A (PKA) activity was measured according to the manufacturer’s instructions (Abcam #ab139435, Cambridge, MA). Stool non-esterified free fatty acid content was measured colorimetrically (HR Series NEFA-HR(2), Wako Diagnostics, Richmond, VA). Indirect calorimetry was performed in individually housed animals over 2 weeks in temperature controlled settings (Columbus Instruments, Columbus, OH), where consumption rates of O_2_ (VO_2_) and CO_2_ (VCO_2_), respiratory exchange ratio (RER), energy expenditure (EE), locomotion (infrared beam breaks), and core body temperature (by intra-abdominal probes, Columbus Instruments, Columbus, OH) were collected simultaneously. Core body temperature was also collected outside of calorimetry cages (SubCue dataloggers, Canadian Analytical Technologies Inc., Calgary, Alberta, Canada). For Scholander plot analysis, mice were acclimated in indirect calorimetry chambers for 2 days before starting the experiment. Cage temperature steps were 10°C, 15°C, 20°C, 25°C, 27°C, 30°C, and 33°C at 2-hour intervals per step. Only data from the second hour at each temperature were used for analysis. For each mouse cohort, Scholander analysis was done at least 3 times consecutively over 3 days and was compiled as averages. For β-guanidino propionic acid (β-GPA) calorimetry experiments, WT and KO mice housed in indirect calorimetry chambers at 30°C and were given vehicle control daily by IP injections. After baseline data were collected for one week, 0.4 g/kg β-GPA was given daily by IP injections and calorimetry data were collected for one week.

### Oral glucose tolerance test

Mice were fasted for 6 hours before injection with 2 g/kg glucose by oral gavage for oral glucose tolerance test (GTT). Blood glucose was measured at 15, 30, 60, 90, and 120 minutes post injection.

### F-18 fluorodeoxyglucose (F-18 FDG) uptake study

Mice were fasted overnight, placed under isoflurane anaesthesia, and given F-18 fluorodeoxyglucose (FDG) (∼0.3mCi/animal) via retro-orbital injection. After 45 minutes, they were sacrificed and iWAT, iBAT, and gastric-soleus muscle were removed and *ex vivo* radioactivity was measured by gamma scintillation counting.

### Inguinal white adipose tissue stromal vascular fraction (SVF) isolation and culture

Isolation and culture of iWAT SVF was performed as previously described ^42^. Briefly, iWAT was removed from mice and digested in Collagenase / Dispase buffer (10mL PBS, 100mg collagenase D, 24mg dispase II, 10mM CaCl_2,_ sterile filtered) for 40 minutes at 37°C and 140 rpm. Digested tissue was filtered through 100uM filter and washed with cold media (DMEM-F12, 10% FBS, 1% penicillin/streptomycin) to inactivate collagenase. Filtered SVF mixture was centrifuged for 10 minutes at 500 x g at 4°C and the supernatant was removed and resuspended in media and filtered through a 70 µM sterile filter. The SVF mixture was again centrifuged at 500 x g at 4°C for 10 minutes. The culture medium was removed and cells were resuspended in fresh medium and plated on collagen coated plates. After 24 hours, plates were washed with PBS to remove debris. SVF were grown to confluence and induced with adipose induction cocktail (0.5 mM IBMX, 1 µM dexamethasone, 850 nM insulin, and 1 µM rosiglitazone) for 48 hours. After 48 hours, medium was switched to contain only 1 µM rosiglitazone and 850 nM insulin. After another 48 hours, medium was switched once again to contain only 850 nM insulin. The SVF culture was completely differentiated by day 7.

### Oil red O staining

Cells were washed with PBS before fixing in 10% phosphate buffered formalin. Cells were washed twice with double distilled H_2_O before incubating with 60% isopropanol for 5 minutes. The cells were then dried completely at room temperature and incubated with Oil Red O for 10 minutes. Oil Red O was then removed and the cells were washed 4 times with double distilled H_2_O before imaging.

### Seahorse

Seahorse assay was performed as previously described ^77^. For tissue oxygen consumption rate, tissue was removed and cut into ∼10-20 mg pieces and placed in 24-well islet capture plates. Tissue was incubated in 750 µL seahorse media (DMEM + Glutamax, 1 mM pyruvate, 25mM glucose) at 37°C until seahorse assay performed within 2 hours of mice sacrifice. 75 µL of 100 µg/ml of oligomycin was injected for a final concentration of 10 µg/ml. The seahorse assay cycles were: mix for 3 minutes, wait for 2 minutes, measure for 3 minutes, repeated 5 times for baseline measurements before injecting with oligomycin. For muscle and iWAT baseline OCR measurements, we adopted previously established protocols as reported in ^78, 79^. Briefly, muscle and iWAT were collected after sacrifice and washed with Krebs-Henseleit buffer (KHB) (111 mM NaCl, 4.7 mM KCl, 2 mM MgSO4, 1.2 mM Na2HPO4, 0.5 mM carnitine, 2.5 mM glucose and 10 mM sodium pyruvate). Tissue were cut into 5-10 mg pieces and plated individually on XF24 islet capture plates. Digitonin was added to permeabilize the membrane. Basal OCR readings were collected with the following cycles: 10 × 2 min measurements, followed by digitonin injection. Subsequent readings were recorded after 2 min mixing and 2 min rest. All OCR values were normalized to individual tissue weights.

### Heart rate and arterial blood pressure

Mixed 129/BL6 mice 2 months old were treated with DMSO vehicle or PV-02076 (20 mg/kg body weight) for two weeks. On the day of the experiment, they were anesthetized with isoflurane and mean arterial blood pressure (MAP) and heart rate were determined over 10 minutes by non-invasive tail cuff (Coda monitor, Kent Scientific Corp, Torrington, CT).

### β-guanidinopropionic acid (β-GPA) SVF seahorse experiments

For β-GPA SVF seahorse experiments, XF24 cell culture microplates were coated with Rat Tail Collagen I (Sigma Cat#C3867) before plating wells with prepared SVF as described previously. On Day 6 of SVF differentiation 50mM of β-GPA or vehicle control were added to the differentiation cocktail. Seahorse assay was run on Day 7. The seahorse assay cycles were: mix for 3 minutes, wait for 2 minutes, measure for 3 minutes, repeated 5 times for baseline measurements. Cells were lysed and protein concentration was measured by DC protein assay (Bio-Rad). OCR values were normalized to protein levels, and baseline OCR values were calculated.

### RNA expression levels

Total adipose tissue RNA was extracted by RNeasy Lipid Tissue Mini Kit (Qiagen), and SVF RNA was extracted with TRIzol (Thermo Fisher) according to manufacturer instructions. RNA expression levels were measured by qRT-PCR with Power SYBR green RNA-to-Ct 1-step kit (thermo fisher/applied biosystems) in 60 ng RNA/10 µL reactions.

### Water repulsion

Water repulsion assay was performed as previously described ^19^. Baseline body temperature was measured rectally by probe thermometer (YSI-73ATA). Mice were allowed to swim in 30° C water for 2 minutes before being placed on a paper towel for a few seconds to remove excess water. Mice were placed in clean cage with no bedding at 22°C, and weight and body temperature were determined every 5 minutes for 60 minutes.

### Thermopreference assay

Thermopreference assay was performed as previously described ^20^. Briefly, three 10-gallon water tanks were used to house mice cages and water heater-circulators were used to maintain water bath temperatures at 22°C, 27°C, and 32°C. Cage temperatures were monitored by thermometer. Cages were connected by translucent tubing to allow freedom of movement across cages. Mouse movement was monitored by a time-lapse, infrared flash overhead camera (Bushnell Model# 119740). Mice were subjected to 5 days of 2-hour per day training on the bench top by connecting two cages with the same tubing. Training multiple mice together increased subsequent multi-cage exploration by single mice in the apparatus. For a given thermopreference assay, one trained mouse was placed into the cages with food and water in all three cages. Mouse data were collected for 4 days at 3 minute time lapse intervals, and time spent in each cage was calculated.

### Acute cold exposure assay

Mice housed at 30°C in individual cages without bedding were brought into a 4°C environment in the same cages for up to 3 hours with ample food and water. Core body temperature was measured every 15 minutes by rectal probe. After the experiment, mice were placed under warm a heat lamp to recover body temperature quickly and were monitored for 1 hour.

### Serum Creatine Kinase Activity assay

To assess muscle activity during cold exposure, we measured serum creatine kinase activity. 100 µl of blood was collected by retro-orbital bleeds before and after acute cold exposure. Blood samples were allowed to coagulate in room temperature for at least 10 minutes before spinning at 5000 x g for 10 minutes. Serum was collected and serum creatine kinase activity was measured according to manufacturer’s instructions (Cat# MAK116, Sigma-Aldrich, St. Louis, MO, USA)

### Gastrointestinal permeability assay

Mouse intestinal permeability was assessed by 4 kDa FITC-Dextran (FD4; Cat# 46944, Sigma-Aldrich, St. Louis, MO, USA) as previously reported ^80^. Briefly, mice were fasted overnight and FD4 was given by oral gavage (0.5mg/g BW). After 90 minutes, plasma was collected by retro-orbital bleeding in EDTA coated tubes (Ref# 365974, Fisher Scientific, Pittsburgh, PA, USA). Plasma was diluted in equal volume PBS and FD4 was measured by fluorometer with an excitation wavelength of 485 nm and emission of 535 nm.

### Immunohistochemistry

Immunohistochemistry staining on adipose tissue were performed by the Albert Einstein College of Medicine Histology and Comparative Pathology Core. Paraffin fixed slides were heated at 60°C for 1 hour before dewaxing (xylene 2 × 10min, 100% ethanol 2×2min, 95% ethanol 2×2 min, 80% ethanol 2×2min, 70% ethanol 2 × 2 min, 70% ethanol 2 × 2 min, water). After dewaxing, slides were washed in TBS buffer twice for 2 minutes each before blocking endogenous peroxidase activity with 3% hydrogen peroxide for 20 minutes at room temperature. Antigen retrieval using 10 mM pH 6.0 Citrate buffer in steamer was performed for 20 minutes, then slides were cooled at room temperature for 30 minutes. Slides were washed again in TBS twice for 3 minutes each before blocking with 2% BSA for 30 minutes at room temperature. Slides were incubated with primary antibody for 60 minutes at room temperature (Tyrosine Hydroxylase Cat# AB75875, Abcam Cambridge, MA, USA, diluted 1:200), then washed 3 times in TBS before applying secondary antibody for 30 minutes at room temperature (Cat# MP-7451, Vector Laboratories, Burlingame, CA, USA). Slides were washed twice for 5 minutes each before applying DAB for 2 minutes. Harris Hematoxylin counterstain was applied for 30 seconds, then the slides were mounted with xylene.

### iWAT Mitochondria Extraction and Citrate Synthase activity assay

Mitochondria were extracted from iWAT using the Mitocheck Mitochondrial Isolation Kit (Cayman Chem Cat# 701010) with a modified protocol. After euthanizing the mice, both iWAT fat pads, with the central lymph node removed, were placed in ice-cold PBS and cut into small pieces. The cut fat pad was transferred into 1 ml of the mitochondrial homogenization buffer and homogenized for 20 seconds in a bead homogenizer (Benchmark Scientific, Edison, NJ). The homogenized solution was centrifuged at 1000 x g for 3 minutes and the supernatant (below the fat layer) was transferred into a fresh tube. The lysate was centrifuged again at 1000 x g for 2 minutes and the supernatant was transferred into another fresh tube to spin at 10,000 x g for 10 minutes. The supernatant was discarded, and the mitochondria pellet was resuspended and washed twice in 1 ml of mitochondrial isolation buffer (10,000 x g for 10 minutes). Finally, the purified mitochondria was resuspended in 50 µl of mitochondrial isolation buffer and kept on ice until used for protein quantification and citrate synthase activity assay according to manufacturer’s instructions (Cayman Chem cat#701040).

## DATA AVAILABILITY

Data that support the findings of this study are available from the corresponding authors on reasonable request.

## ACKNOWLEDGMENTS

We thank Barbara Cannon and Jan Nedergaard for helpful discussions. This work was supported by NIH grants GM007491 (VJP), DK020541 (VLS), DK10541 and DK020541 (GJS), and AG043517 and AG031782 (RS); by American Diabetes Association grant 1-18-IBS-062 (RJ); and by the Harrington Discovery Institute (VLS). At Albert Einstein College of Medicine, we thank the Gruss Magnetic Resonance Research Center, the Analytical Imaging Facility, the Histotechnology and Comparative Pathology Facility, the Einstein Norman Fleischer Diabetes Center, and Xue-liang Du of the Seahorse Assay facility. Creatinine measurements were performed by the University of Alabama - UCSD O’Brien Kidney Center NIH DK079337.

## AUTHOR CONTRIBUTIONS

V.J.P., R.S, G.J.S., and V.L.S. conceptualised the study. V.J.P., R.L., Y.C., and V.L.S. provided methodology. V.J.P., L.W., M.G.M., and V.L.S. provided format analysis. V.J.P., R.L., L.W., M.G.M., W.R.K., and Y.C. performed investigations. V.J.P., L.W., W.R.K., Y.C., R. S., G.J.S., and V.L.S. provided resources. V.J.P. and V.L.S. wrote the original draft. V.J.P., R.S., G.J.S., and V.L.S. were involved in review and editing. V.J.P. and V.L.S. provided study visualization. V.L.S. provided study supervision and administration. R.S., G.J.S., V.L.S. provided funding acquisition.

## REFERENCES

1 Lipinski, B. A. & Mathias, M. M. Prostaglandin production and lipolysis in isolated rat adipocytes as affected by dietary fat. Prostaglandins 16, 957–963 (1978).

2 Michaud, A. et al. Prostaglandin (PG) F2 alpha synthesis in human subcutaneous and omental adipose tissue: modulation by inflammatory cytokines and role of the human aldose reductase AKR1B1. PloS one 9, e90861 (2014).

3 Pisani, D. F. et al. The ω6-fatty acid, arachidonic acid, regulates the conversion of white to brite adipocyte through a prostaglandin/calcium mediated pathway. Molecular metabolism 3, 834–847 (2014).

4 Kanai, N. et al. Identification and characterization of a prostaglandin transporter. Science 268, 866–869 (1995).

5 Schuster, V. L. Molecular mechanisms of prostaglandin transport. Annu Rev Physiol 60, 221–242 (1998).

6 Nomura, T., Lu, R., Pucci, M. L. & Schuster, V. L. The two-step model of prostaglandin signal termination: in vitro reconstitution with the prostaglandin transporter and prostaglandin 15 dehydrogenase. Mol Pharmacol 65, 973–978 (2004).

7 Chi, Y., Suadicani, S. O. & Schuster, V. L. Regulation of prostaglandin EP1 and EP4 receptor signaling by carrier-mediated ligand reuptake. Pharmacology research & perspectives 2, e00051, doi:10.1002/prp2.51 (2014).

8 Reginato, M. J., Krakow, S. L., Bailey, S. T. & Lazar, M. A. Prostaglandins promote and block adipogenesis through opposing effects on peroxisome proliferator-activated receptor gamma. J Biol Chem 273, 1855–1858 (1998).

9 Taketani, Y. et al. Activation of the prostanoid FP receptor inhibits adipogenesis leading to deepening of the upper eyelid sulcus in prostaglandin-associated periorbitopathy. Investigative ophthalmology & visual science 55, 1269–1276, doi:10.1167/iovs.13-12589 (2014).

10 Volat, F. E. et al. Depressed levels of prostaglandin F2alpha in mice lacking Akr1b7 increase basal adiposity and predispose to diet-induced obesity. Diabetes 61, 2796–2806, doi:10.2337/db11-1297 (2012).

11 Vegiopoulos, A. et al. Cyclooxygenase-2 controls energy homeostasis in mice by de novo recruitment of brown adipocytes. Science 328, 1158–1161, doi:10.1126/science.1186034 (2010).

12 Madsen, L. et al. UCP1 induction during recruitment of brown adipocytes in white adipose tissue is dependent on cyclooxygenase activity. PLoS One 5, e11391, doi:10.1371/journal.pone.0011391 (2010).

13 García-Alonso, V. et al. Prostaglandin E 2 Exerts Multiple Regulatory Actions on Human Obese Adipose Tissue Remodeling, Inflammation, Adaptive Thermogenesis and Lipolysis. PloS one 11, e0153751 (2016).

14 Garcia-Alonso, V. et al. Coordinate functional regulation between microsomal prostaglandin E synthase-1 (mPGES-1) and peroxisome proliferator-activated receptor gamma (PPARgamma) in the conversion of white-to-brown adipocytes. J Biol Chem 288, 28230–28242, doi:10.1074/jbc.M113.468603 (2013).

15 Kondepudi, D. Introduction to modern thermodynamics. (Wiley, 2008).

16 Halford, J. C., Wanninayake, S. C. & Blundell, J. E. Behavioral satiety sequence (BSS) for the diagnosis of drug action on food intake. Pharmacology Biochemistry and Behavior 61, 159–168 (1998).

17 Virtue, S., Even, P. & Vidal-Puig, A. Below thermoneutrality, changes in activity do not drive changes in total daily energy expenditure between groups of mice. Cell metabolism 16, 665–671 (2012).

18 Nedergaard, J. & Cannon, B. The browning of white adipose tissue: some burning issues. Cell Metab 20, 396–407, doi:10.1016/j.cmet.2014.07.005 (2014).

19 Westerberg, R. et al. Role for ELOVL3 and fatty acid chain length in development of hair and skin function. Journal of Biological Chemistry 279, 5621–5629 (2004).

20 Gaskill, B. N., Rohr, S. A., Pajor, E. A., Lucas, J. R. & Garner, J. P. Working with what you’ve got: Changes in thermal preference and behavior in mice with or without nesting material. Journal of Thermal Biology 36, 193–199 (2011).

21 Abreu-Vieira, G., Xiao, C., Gavrilova, O. & Reitman, M. L. Integration of body temperature into the analysis of energy expenditure in the mouse. Molecular metabolism 4, 461–470 (2015).

22 Scholander, P., Hock, R., Walters, V. & Irving, L. Adaptation to cold in arctic and tropical mammals and birds in relation to body temperature, insulation, and basal metabolic rate. The Biological Bulletin 99, 259–271 (1950).

23 Fischer, A. W., Csikasz, R. I., von Essen, G., Cannon, B. & Nedergaard, J. No insulating effect of obesity. American Journal of Physiology-Endocrinology and Metabolism 311, E202–E213 (2016).

24 Kazak, L. et al. A Creatine-Driven Substrate Cycle Enhances Energy Expenditure and Thermogenesis in Beige Fat. Cell 163, 643–655 (2015).

25 Feldmann, H. M., Golozoubova, V., Cannon, B. & Nedergaard, J. UCP1 ablation induces obesity and abolishes diet-induced thermogenesis in mice exempt from thermal stress by living at thermoneutrality. Cell metabolism 9, 203–209 (2009).

26 Chouchani, E. T., Kazak, L. & Spiegelman, B. M. New Advances in Adaptive Thermogenesis: UCP1 and Beyond. Cell Metabolism 29, 27–37 (2018).

27 Schuster, V. L. & Chi, Y. (Google Patents, 2017).

28 Chi, Y. et al. Development of a high-affinity inhibitor of the prostaglandin transporter. J Pharmacol Exp Ther 339, 633–641, doi:10.1124/jpet.111.181354 (2011).

29 Golozoubova, V. et al. Only UCP1 can mediate adaptive nonshivering thermogenesis in the cold. The FASEB Journal 15, 2048–2050 (2001).

30 Enerback, S. et al. Mice lacking mitochondrial uncoupling protein are cold-sensitive but not obese. Nature 387, 90–93 (1997).

31 Molderings, G. J., Likungu, J. & Göthert, M. Modulation of noradrenaline release from the sympathetic nerves of human right atrial appendages by presynaptic EP3-and DP-receptors. Naunyn-Schmiedeberg’s archives of pharmacology 358, 440–444 (1998).

32 Exner, H. J. & Schlicker, E. Prostanoid receptors of the EP3 subtype mediate the inhibitory effect of prostaglandin E2 on noradrenaline release in the mouse brain cortex. Naunyn-Schmiedeberg’s archives of pharmacology 351, 46–52 (1995).

33 Molderings, G., Malinowska, B. & Schlicker, E. Inhibition of noradrenaline release in the rat vena cava via prostanoid receptors of the EP3-subtype. British journal of pharmacology 107, 352–355 (1992).

34 Baines, A. D. Effects of salt intake and renal denervation on catecholamine catabolism and excretion. Kidney international 21, 316–322 (1982).

35 Lepschy, M., Rettenbacher, S., Touma, C. & Palme, R. Excretion of catecholamines in rats, mice and chicken. Journal of Comparative Physiology B 178, 629–636 (2008).

36 Esler, M. et al. Overflow of catecholamine neurotransmitters to the circulation: source, fate, and functions. Physiol Rev 70, 963–985 (1990).

37 Chi, Y. et al. Inhibition of the Prostaglandin Transporter PGT Lowers Blood Pressure in Hypertensive Rats and Mice. PloS one 10, e0131735 (2015).

38 Gehrmann, J. et al. Phenotypic screening for heart rate variability in the mouse. American Journal of Physiology-Heart and Circulatory Physiology 279, H733–H740 (2000).

39 Ishii, K., Kuwahara, M., Tsubone, H. & Sugano, S. Autonomic nervous function in mice and voles (Microtus arvalis): investigation by power spectral analysis of heart rate variability. Laboratory animals 30, 359–364 (1996).

40 Contreras, G. A., Lee, Y.-H., Mottillo, E. P. & Granneman, J. G. Inducible brown adipocytes in subcutaneous inguinal white fat: the role of continuous sympathetic stimulation. American Journal of Physiology-Endocrinology and Metabolism 307, E793–E799 (2014).

41 Scott, M. A., Nguyen, V. T., Levi, B. & James, A. W. Current methods of adipogenic differentiation of mesenchymal stem cells. Stem cells and development 20, 1793–1804 (2011).

42 Rodbell, M. Metabolism of isolated fat cells I. Effects of hormones on glucose metabolism and lipolysis. J Biol Chem 239, 375–380 (1964).

43 Sears, I. B., MacGinnitie, M. A., Kovacs, L. G. & Graves, R. A. Differentiation-dependent expression of the brown adipocyte uncoupling protein gene: regulation by peroxisome proliferator-activated receptor gamma. Molecular and cellular biology 16, 3410–3419 (1996).

44 Jones, J. R. et al. Deletion of PPARγ in adipose tissues of mice protects against high fat diet-induced obesity and insulin resistance. Proceedings of the National Academy of Sciences 102, 6207–6212 (2005).

45 Xu, L. et al. Ablation of PPAR γ in subcutaneous fat exacerbates age-associated obesity and metabolic decline. Aging cell 17, e12721 (2018).

46 Klepac, K. et al. The Gq signalling pathway inhibits brown and beige adipose tissue. Nature communications 7, 1–10 (2016).

47 Griffin, B. W., Klimko, P., Crider, J. Y. & Sharif, N. A. AL-8810: a novel prostaglandin F2 alpha analog with selective antagonist effects at the prostaglandin F2 alpha (FP) receptor. J Pharmacol Exp Ther 290, 1278–1284 (1999).

48 Glushakov, A. V., Robbins, S. W., Bracy, C. L., Narumiya, S. & Doré, S. Prostaglandin F 2α FP receptor antagonist improves outcomes after experimental traumatic brain injury. Journal of neuroinflammation 10, 132 (2013).

49 Kim, Y. T., Moon, S. K., Maruyama, T., Narumiya, S. & Doré, S. Prostaglandin FP receptor inhibitor reduces ischemic brain damage and neurotoxicity. Neurobiology of disease 48, 58–65 (2012).

50 Ukropec, J., Anunciado, R. P., Ravussin, Y., Hulver, M. W. & Kozak, L. P. UCP1-independent thermogenesis in white adipose tissue of cold-acclimated Ucp1-/- mice. Journal of Biological Chemistry 281, 31894–31908 (2006).

51 Serrero, G., Lepak, N. M. & Goodrich, S. P. Prostaglandin F2α inhibits the differentiation of adipocyte precursors in primary culture. Biochemical and biophysical research communications 183, 438–442 (1992).

52 Lepak, N. & Serrero, G. Inhibition of adipose differentiation by 9α, 11β-prostaglandin F2α. Prostaglandins 46, 511–517 (1993).

53 Casimir, D. A., Miller, C. W. & Ntambi, J. M. Preadipocyte differentiation blocked by prostaglandin stimulation of prostanoid FP2 receptor in murine 3T3-L1 cells. Differentiation; research in biological diversity 60, 203–210, doi:10.1046/j.1432-0436.1996.6040203.x (1996).

54 Liu, L. & Clipstone, N. A. Prostaglandin F2α inhibits adipocyte differentiation via a Gαq-Calcium-Calcineurin-Dependent signaling pathway. Journal of cellular biochemistry 100, 161–173 (2007).

55 Annamalai, D. & Clipstone, N. A. Prostaglandin F2α inhibits adipogenesis via an autocrine-mediated interleukin-11/glycoprotein 130/STAT1-dependent signaling cascade. Journal of cellular biochemistry 115, 1308–1321 (2014).

56 Shabalina, I. G. et al. UCP1 in brite/beige adipose tissue mitochondria is functionally thermogenic. Cell reports 5, 1196–1203 (2013).

57 Stawski, C., Koteja, P. & Sadowska, E. T. A Shift in the Thermoregulatory Curve as a Result of Selection for High Activity-Related Aerobic Metabolism. Front Physiol 8, 1070, doi:10.3389/fphys.2017.01070 (2017).

58 Stawski, C., Koteja, P., Sadowska, E. T., Jefimow, M. & Wojciechowski, M. S. Selection for high activity-related aerobic metabolism does not alter the capacity of non-shivering thermogenesis in bank voles. Comparative Biochemistry and Physiology Part A: Molecular & Integrative Physiology 180, 51–56 (2015).

59 Golozoubova, V., Cannon, B. & Nedergaard, J. UCP1 is essential for adaptive adrenergic nonshivering thermogenesis. American Journal of Physiology-Endocrinology and Metabolism 291, E350–E357 (2006).

60 Hofmann, W. E., Liu, X., Bearden, C. M., Harper, M.-E. & Kozak, L. P. Effects of genetic background on thermoregulation and fatty acid-induced uncoupling of mitochondria in UCP1-deficient mice. Journal of Biological Chemistry 276, 12460–12465 (2001).

61 Pollard, A. E. et al. AMPK activation protects against diet induced obesity through Ucp1-independent thermogenesis in subcutaneous white adipose tissue. Nat Metab 1, 340–349, doi:10.1038/s42255-019-0036-9 (2019).

62 Danneskiold-Samsøe, N. B. et al. Overexpression of cyclooxygenase-2 in adipocytes reduces fat accumulation in inguinal white adipose tissue and hepatic steatosis in high-fat fed mice. Scientific Reports 9, 8979 (2019).

63 Burl, R. B. et al. Deconstructing adipogenesis induced by β3-adrenergic receptor activation with single-cell expression profiling. Cell metabolism 28, 300–309 (2018).

64 Kazak, L. et al. Genetic Depletion of Adipocyte Creatine Metabolism Inhibits Diet-Induced Thermogenesis and Drives Obesity. Cell Metab 26, 660–671, doi:10.1016/j.cmet.2017.08.009 (2017).

65 Kazak, L. et al. Ablation of adipocyte creatine transport impairs thermogenesis and causes diet-induced obesity. Nature Metabolism 1, 360–370 (2019).

66 Cheng, Y. et al. Prediction of Adipose Browning Capacity by Systematic Integration of Transcriptional Profiles. Cell reports 23, 3112–3125 (2018).

67 Shan, T. et al. Distinct populations of adipogenic and myogenic Myf5-lineage progenitors in white adipose tissues. Journal of lipid research 54, 2214–2224 (2013).

68 Sanchez-Gurmaches, J. et al. PTEN loss in the Myf5 lineage redistributes body fat and reveals subsets of white adipocytes that arise from Myf5 precursors. Cell metabolism 16, 348–362 (2012).

69 Ikeda, K. et al. UCP1-independent signaling involving SERCA2b-mediated calcium cycling regulates beige fat thermogenesis and systemic glucose homeostasis. Nat Med 23, 1454–1465, doi:10.1038/nm.4429 (2017).

70 Bertholet, A. M. et al. Mitochondrial patch clamp of beige adipocytes reveals UCP1-positive and UCP1-negative cells both exhibiting futile creatine cycling. Cell metabolism 25, 811–822 (2017).

71 Chouchani, E. T. & Kajimura, S. Metabolic adaptation and maladaptation in adipose tissue. Nature Metabolism 1, 189–200, doi:10.1038/s42255-018-0021-8 (2019).

72 Mottillo, E. P. et al. Coupling of lipolysis and de novo lipogenesis in brown, beige, and white adipose tissues during chronic β3-adrenergic receptor activation. Journal of lipid research 55, 2276–2286 (2014).

73 Marlatt, K. L., Chen, K. Y. & Ravussin, E. Is Activation of Human Brown Adipose Tissue a Viable Target for Weight Management? American journal of physiology. Regulatory, integrative and comparative physiology 315, R479–R483 (2018).

74 Anunciado-Koza, R., Ukropec, J., Koza, R. A. & Kozak, L. P. Inactivation of UCP1 and the glycerol phosphate cycle synergistically increases energy expenditure to resist diet-induced obesity. Journal of Biological Chemistry 283, 27688–27697 (2008).

75 Riachi, M., Himms-Hagen, J. & Harper, M.-E. Percent relative cumulative frequency analysis in indirect calorimetry: application to studies of transgenic mice. Canadian journal of physiology and pharmacology 82, 1075–1083 (2004).

76 Chang, H. Y., Locker, J., Lu, R. & Schuster, V. L. Failure of postnatal ductus arteriosus closure in prostaglandin transporter-deficient mice. Circulation 121, 529–536, doi:CIRCULATIONAHA.109.862946 [pii] 10.1161/CIRCULATIONAHA.109.862946 (2010).

77 Bugge, A., Dib, L. & Collins, S. in Methods Enzymol Vol. 538 233–247 (Elsevier, 2014).

78 Martinez-Lopez, N. et al. Autophagy in the CNS and Periphery Coordinate Lipophagy and Lipolysis in the Brown Adipose Tissue and Liver. Cell metabolism 23, 113–127 (2016).

79 Martinez-Lopez, N. et al. System-wide Benefits of Intermeal Fasting by Autophagy. Cell Metab 26, 856–871.e855, doi:10.1016/j.cmet.2017.09.020 (2017).

80 Dong, C. X. et al. The intestinal epithelial insulin-like growth factor-1 receptor links glucagon-like peptide-2 action to gut barrier function. Endocrinology 155, 370–379 (2014).

