## Supplemental Figures for "Inhibiting the prostaglandin transporter PGT induces non-canonical thermogenesis at thermoneutrality"

***Supplementary Figure 1.***

***Metabolic phenotyping of PGT-KO mice housed at 22°C eating normal chow.*** **(A)** PGT gene expression among different fat pads by qRT-PCR, n=4. **(B)** Decreased fat pad weights in PGT-KO mice, n=6 per group. **(C)** Area under curve of glucose tolerance test of Figure 1P. **(D-H)** Decreased leptin gene expression in PGT-KO mice, as measured by qRT-PCR, n=6 per group; serum levels of leptin, insulin, free fatty acids, and adiponectin in WT and PGT-KO mice, n=4 per group. **(I-K)** Representative iWAT H&E stains of WT, PGT heterozygous, and PGT-KO mice, bar = 50 µm. **(L)** Decreased average adipocyte size in PGT-KO iWAT, n=4 mouse per group. **(M)** Representative H&E stain of multilocular adipocytes in iWAT of PGT-KO mice, bar = 10 µm. **(N-O)** Representative H&E stains of iBAT from WT and PGT-KO mice, bar = 50 µm. Values are mean ± SEM. (*$P<0.05$, **$P<0.01$, ***$P<0.001$, versus respective control; Student's t-test).

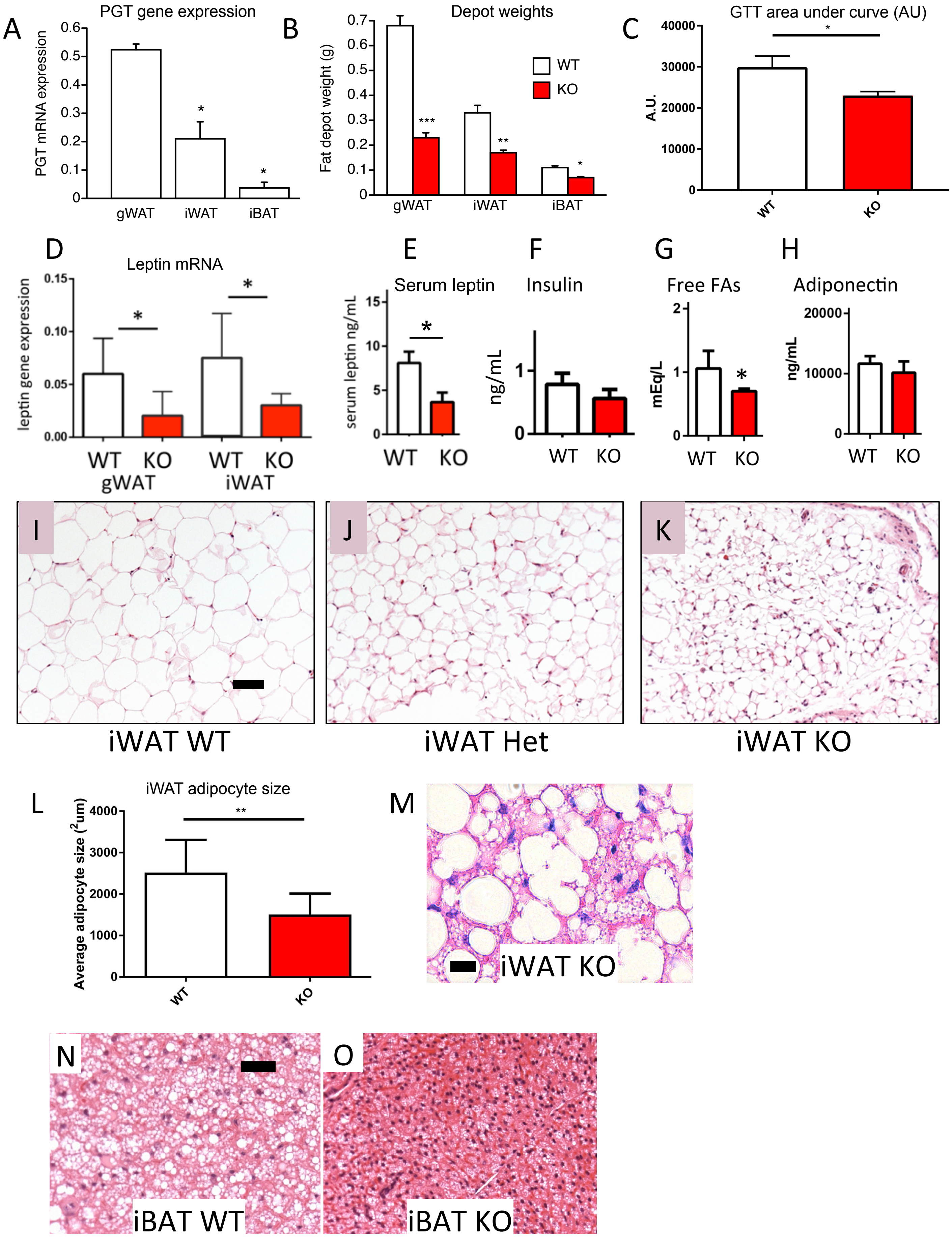


Pai et al Supplementary Figure 1

***Supplementary Figure 2.***

***Phenotypic changes in PGT-KO mice.*** **(A)** Food intake per gram body weight. **(B)** No difference in intestinal integrity between WT and PGT-KO mice as measured by stool weight, stool free fatty acid, and intestinal permeability (as determined by permeability to FITC-4kDA dextran). **(C)** No difference in gastroc-soleus muscle mitochondrial gene expression markers (cox5b and PGC1α) and baseline muscle oxygen consumption rate. **(D)** Representative photo of excised WT and PGT-KO iWAT showing significant browning of the latter. **(E)** No difference in liver citrate synthase activity between WT and PGT-KO mice.

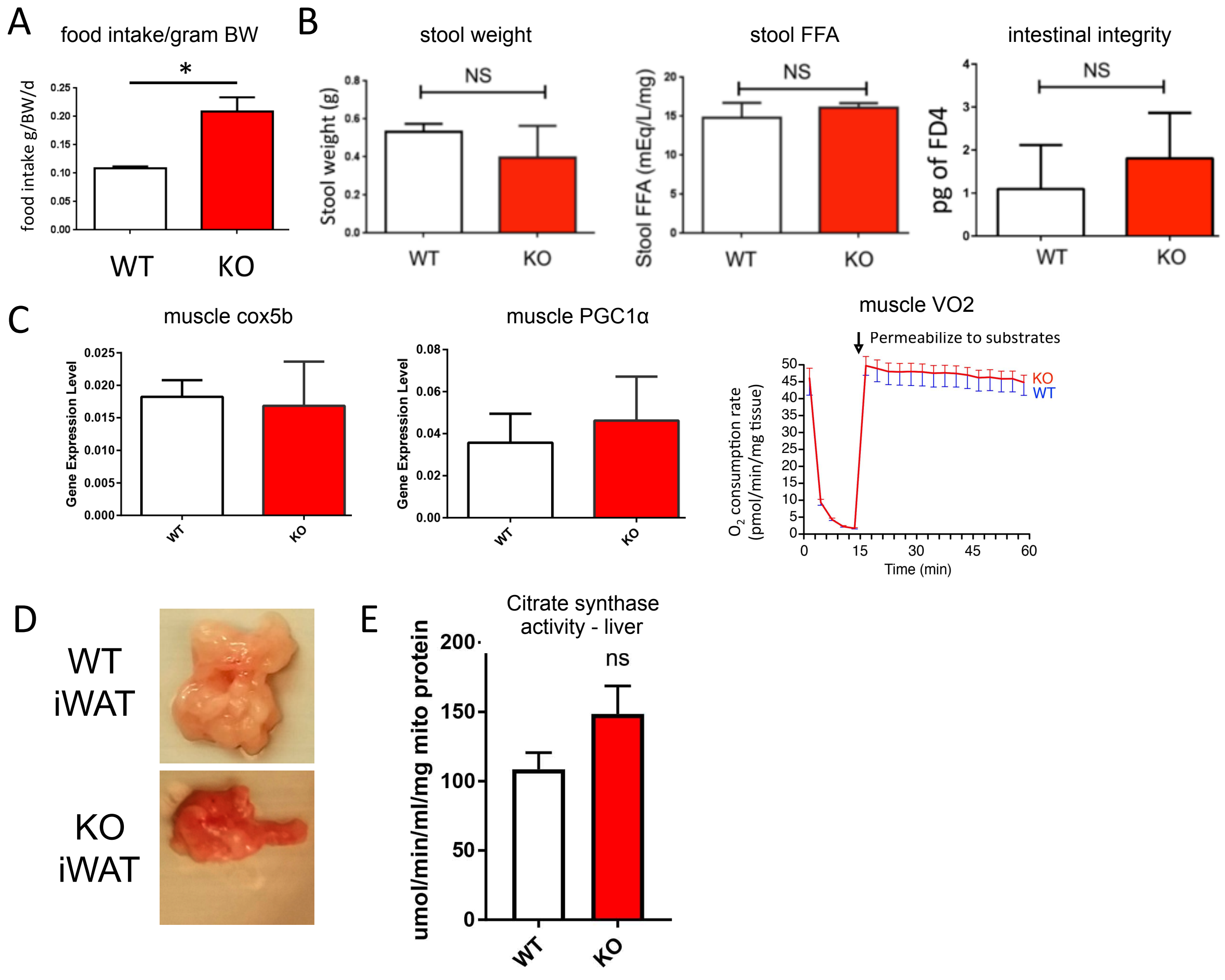


Pai et al Supplementary Figure 2

***Supplementary Figure 3***.

***Primary thermogenesis in PGT-KO mice.*** **(A)** Normal skin and tail in PGT-KO mice as assessed by water repulsion assay, n=4 per group. **(B)** Schematic of thermopreference assay. **(C)** Heat loss induces change in thermopreference, as measured by shaving fur off C57BL/6 mice. Paired experimental design, n=4. **(D-E)** Increased $VO_2$ in PGT-KO mice housed at thermoneutrality, but no significant difference in respiratory exchange ratio (RER), n=4 per group. (**F)** Improved defense against acute cold exposure (4˚C) without shivering in PGT-KO mouse, using serum creatine kinase activity as a shivering index (Kazak et al., 2015, reference #24), n=4 per group. **(G)** Body composition analysis of WT and PGT-KO mice housed at thermoneutrality for 1 month and fed 60% high fat diet, n=8 per group. **(H)** Increased oxygen consumption rate per lean body weight in WT and PGT-KO mice fed high fat diet and housed at 30˚C. **(I)** Increased oxygen consumption rate per mg tissue in iWAT explants from PGT-KO mice fed high fat diet and housed at 30˚C. Digitonin is used at arrow time point to permeabilize substrates, n=10 per group. **(J)** Increased whole fat pad oxygen consumption rate in PGT-KO mice. Data normalized to total fat pad weight from (I). **(K)** Calculated area under curve for iWAT VO2, n=10 per group. **(L)** Increase in browning gene expression in PGT-KO iWAT (right 2 bars in each panel); gWAT gene expression is unchanged (left 2 bars in each panel), n=8 per group. Values are mean ± SEM. (*P<0.05, **P<0.01, versus respective control; Student's t-test).

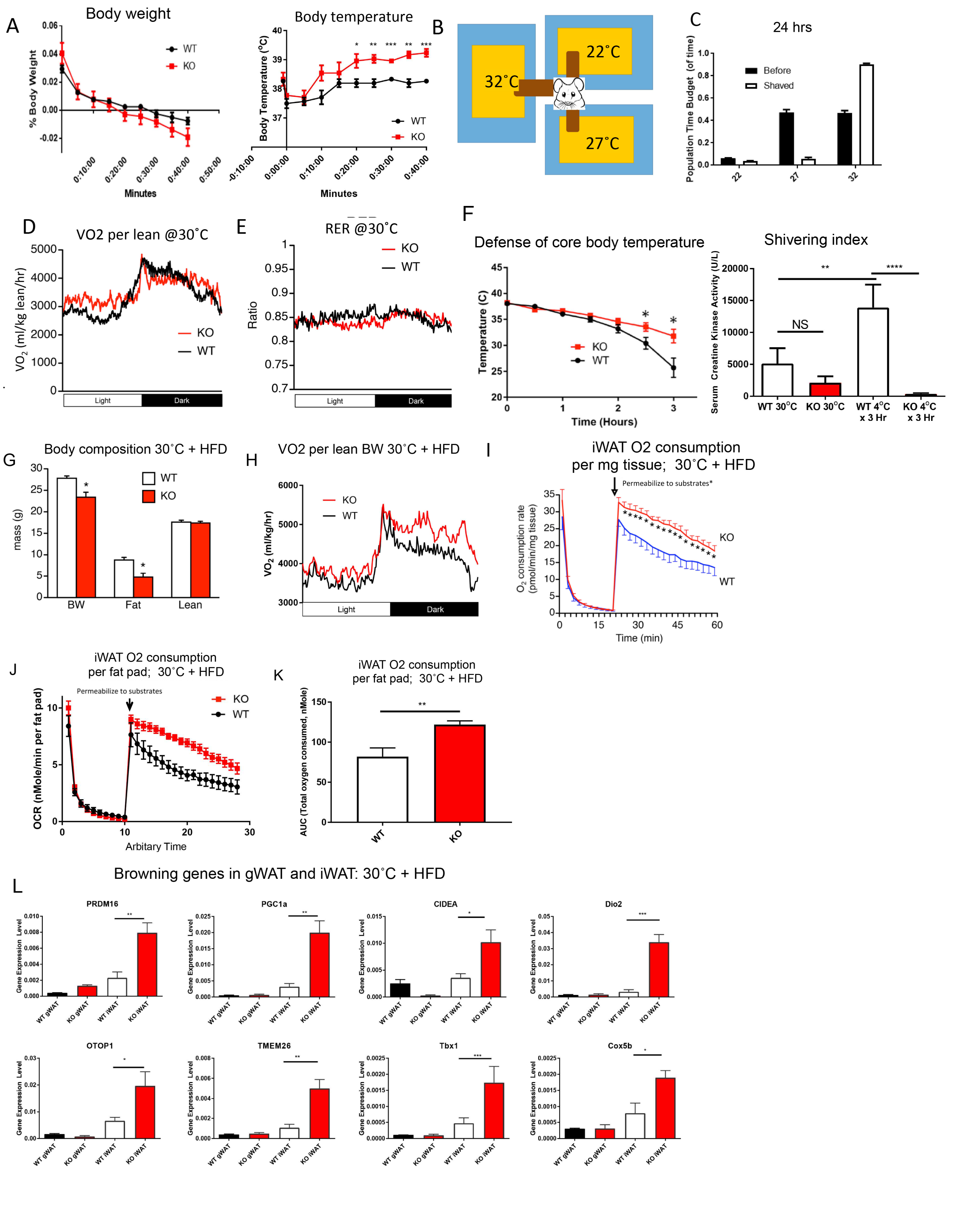


Pai et al Supplementary Figure 3

***Supplementary Figure 4.***
***Treatment of C57BL/6 mice with high affinity PGT inhibitor PV-02076 daily and fed 60% high fat diet recapitulates thermogenesis phenotype of PGT-KO mice.*** **(A)** Increased urinary $PGE_2$ and $PGF_{2\alpha}$ in mice administered PV-02076 (PV) as compared to vehicle control (DMSO). **(B)** No difference in food intake between mice given PV-02076 and vehicle control mice. **(C)** Decreased total fat mass in mice given PV-02076 as measured by echo-MRI. **(D)** Decrease in total body weight gain in mice given PV-02076 as compared to DMSO control when fed 60% high fat diet. **(E)** Increased $VO_2$ in mice given PV-02076 as measured by indirect calorimetry. **(F)** Improved glucose tolerance test in mice given PV-02076. **(G)** Induction of beige fat markers Dio2 and Cidea by PV-02076 in iWAT of C57BL/6 mice.  n=6 per group. Values are mean ± SEM. (*P<0.05, versus respective control; Student's t-test).

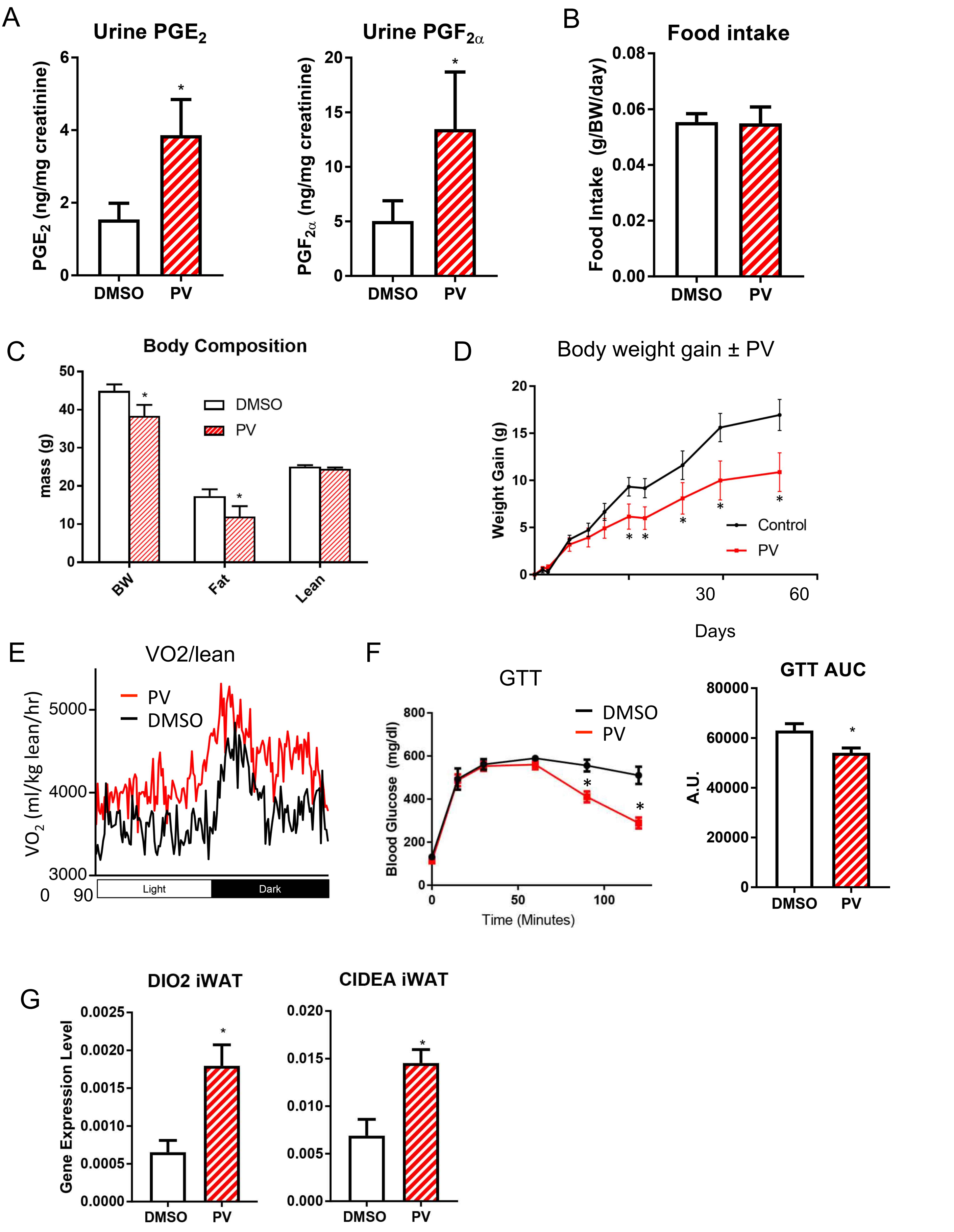


Pai et al Supplementary Figure 4

***Supplementary Figure 5.***

***Suppression of UCP1 in the PGT-KO mice.*** **(A)** Lack of induction of UCP1 in iWAT and iBAT in mice housed at 30˚C and 22˚C, and after exposure to 4˚C for 16 hours, depicted as raw CT values of qRT-PCR n=4 per group where lower Ct value means higher gene expression. **(B)** Heart rate, systolic blood pressure, and diastolic blood pressure in 129/BL6 mice treated for 2 weeks with either vehicle (DMSO) or PV-02075 20 mg/kg BW intraperitoneally. n = 10 each. **(C)** No difference in urinary (systemic) epinephrine levels between WT and PGT-KO mice. n=4 per group. **(D)** Representative photomicrographs of immunohistochemistry stain of tyrosine kinase in iWAT of WT and PGT-KO mice. Bar = 10 µm. Values are mean ± SEM. (*P<0.05, versus respective control; Student's t-test).

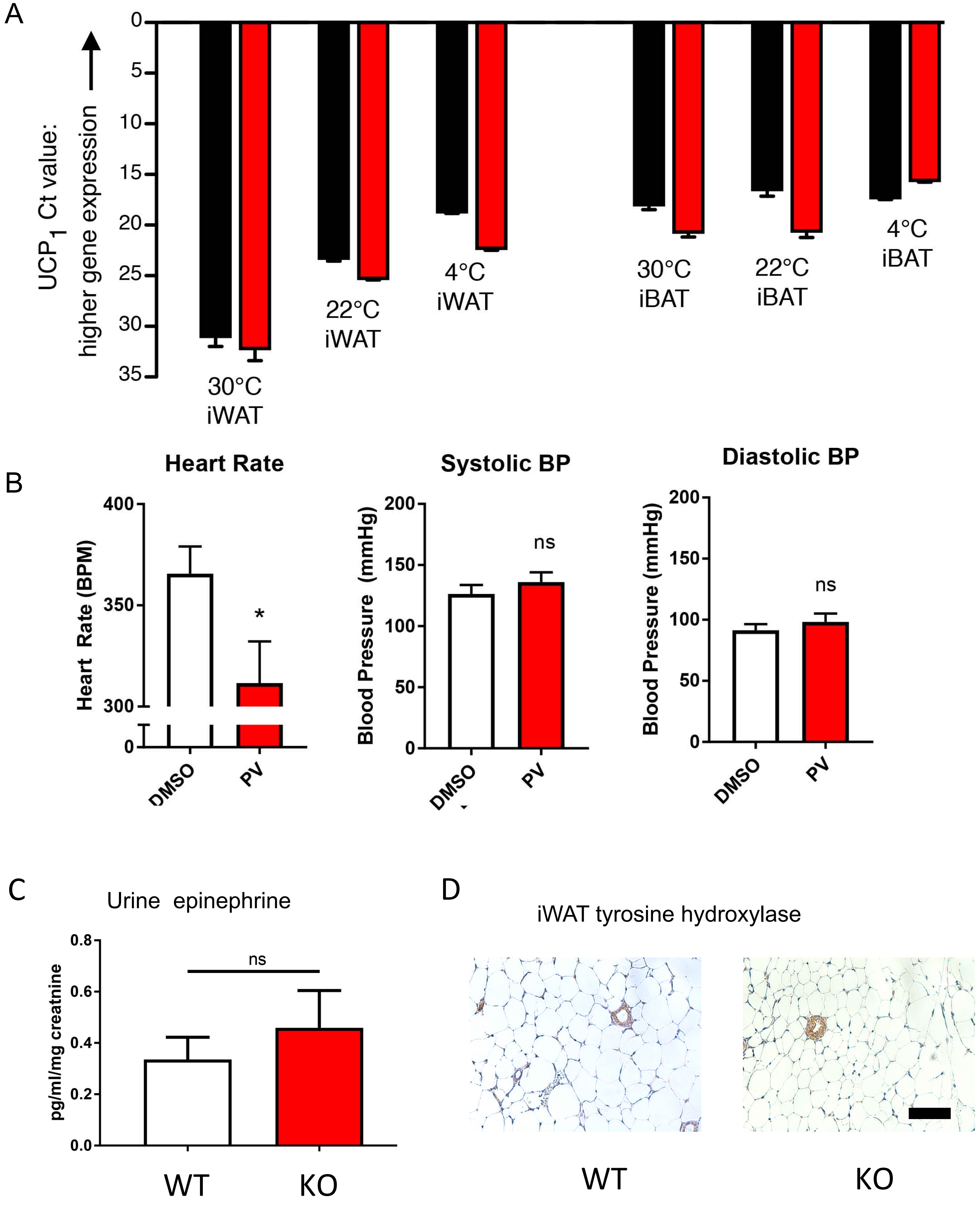


Pai et al Supplementary Figure 5

***Supplementary Figure 6.***
***Evidence supporting induction of futile cycle in PGT-KO mice.***
**(A)** No difference in $VO_2$ of PGT-KO mice administered the creatine transporter *Slc6a8* inhibitor β-guanidinopropionic acid (β-GPA) for one week and housed at thermoneutrality. **(B)** Percent relative cumulative frequency analysis of VO2 data in (A), n=4 per group. **(C)** Microarray fold change intensity ratio plot between WT and PGT-KO iWAT: ATP2a1 (Serca1) indicated as shown. Increase in ATP2a1 gene expression level confirmed in thermoneutral mouse PGT-KO iWAT by qRT-PCR, n=8 per group. **(D)** Induction of Myf5 gene expression in iWAT of PGT-KO mice by qRT-PCR. **(E)** Increase in fatty β-oxidation gene expression in PGT-KO iWAT, measured by qRT-PCR. **(F)** No change in iWAT lipolysis genes. **(G)** No change in iWAT lipogenesis genes. n=8 per group. Values are mean ± SEM. (*P<0.05, **P<0.01, versus respective control; Student's t-test).
<u>Fatty β-oxidation gene names</u>: Acadvl = very long-chain acyl-CoA dehydrogenase; Acaa2 = acetyl-Coenzyme A acyltransferase 2; Cpt1b = carnitine palmitoyltransferase 1B; Cpt2 = carnitine palmitoyltransferase 2; Acads = Acyl-CoA dehydrogenase short chain; Acadl = Acyl-CoA dehydrogenase long chain
<u>Lipolysis gene names</u>: ATGL = adipose triglyceride lipase; HSL = hormone-sensitive lipase
<u>Lipogenesis gene names</u>: SREBP-1c = sterol regulatory element binding protein-1c; SCD1 = steroyl CoA desaturase-1; FASN = fatty acid synthase.

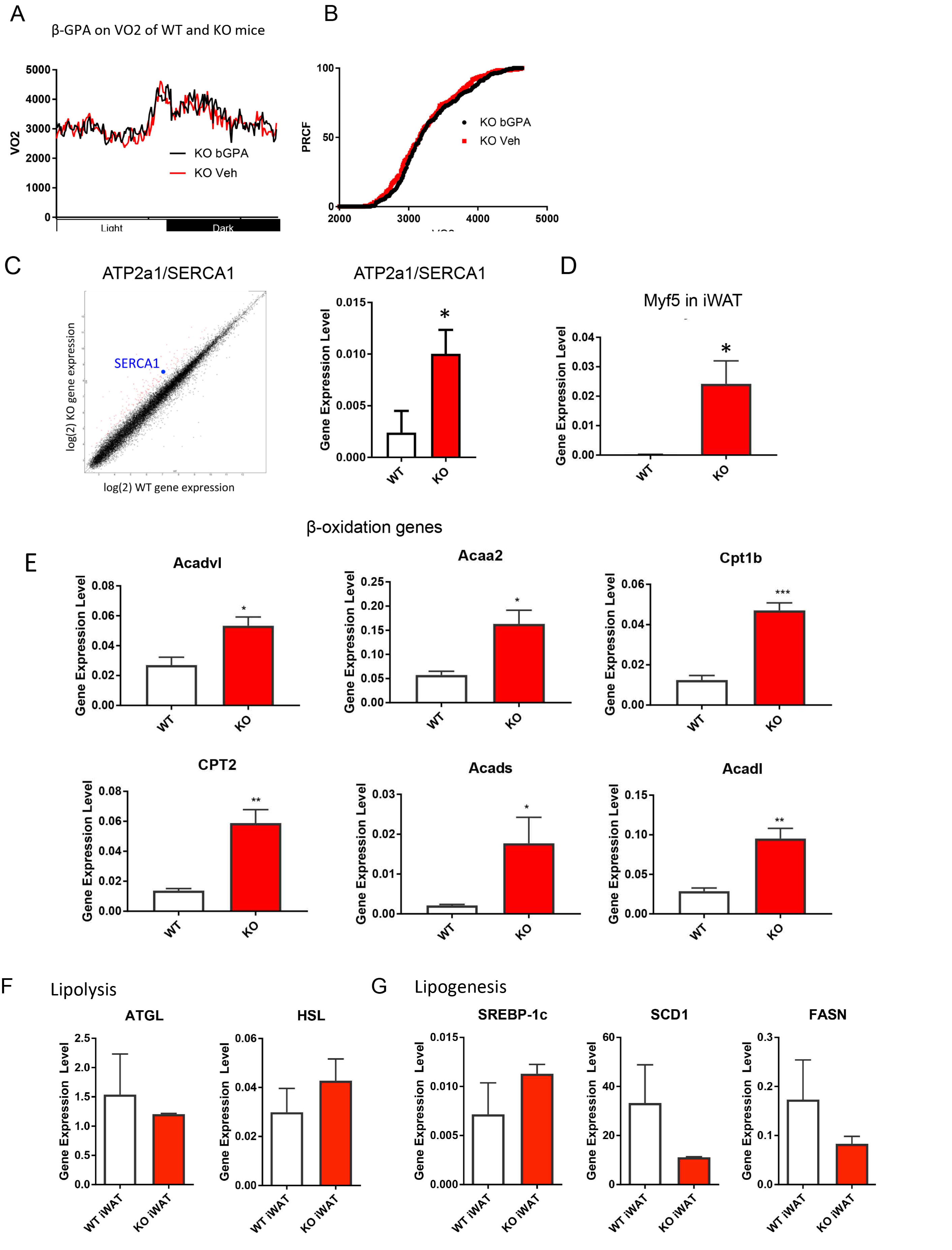


Pai et al Supplementary Figure 6

**Supplementary Figure 7.**

***Proposed model for non-canonical thermogenesis at thermoneutrality in PGT-KO mice.*** While increased $PGE_2$ in PGT-KO mice inhibits facultative norepinephrine release from sympathetic nerve terminals, $PGE_2$ can still activate cAMP signaling constitutively to induce PGC1α activation and mitochondrial biogenesis. Increased β oxidation of fatty acids drives increased ATP synthesis by the expanded mitochondrial pool. The accompanying increase in $PGF_{2\alpha}$ in PGT-KO mice activates the receptor FP which, via Gαq signaling, reduces UCP1 gene expression by inhibiting PPARγ gene expression and induces components of the creatine shuttle. The increased ATP synthesis, via the creatine shuttle, supports UCP1-independent thermogenesis via (unidentified) futile cycle(s).

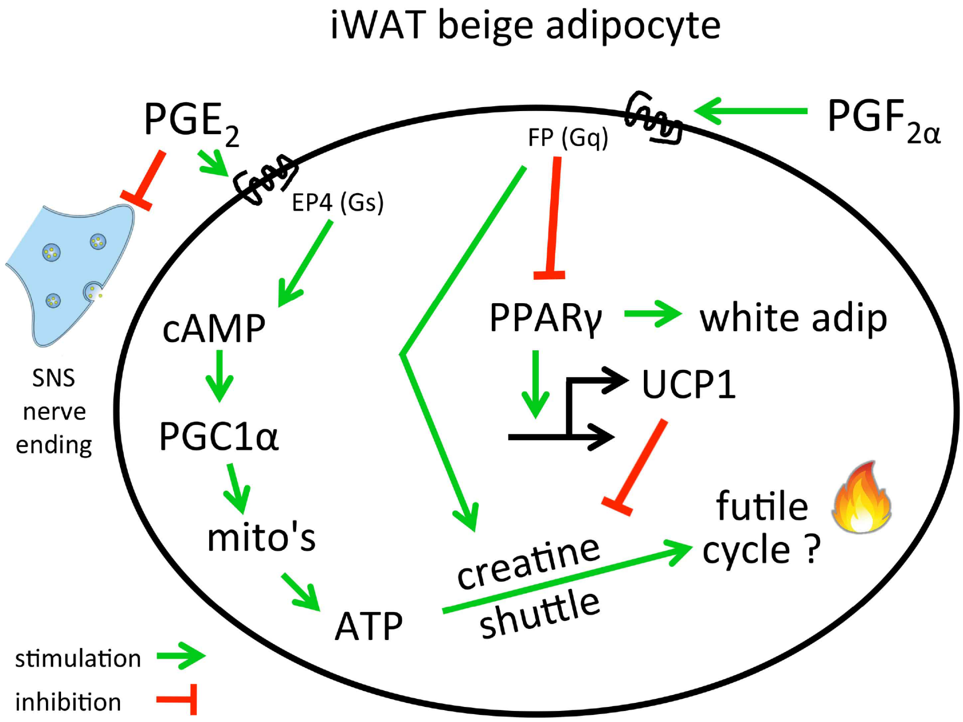
